# Identification of new MmpL3 inhibitors by untargeted and targeted mutant screens defines MmpL3 domains with differential resistance

**DOI:** 10.1101/564245

**Authors:** John T. Williams, Elizabeth R. Haiderer, Garry B. Coulson, Kayla N. Conner, Edmund Ellsworth, Chao Chen, Thomas Dick, Robert B. Abramovitch

## Abstract

The *Mycobacterium tuberculosis* (Mtb) mycolic acid flippase MmpL3 has been the proposed target for multiple inhibitors with diverse chemical scaffolds. This diversity in chemical scaffolds has made it difficult to predict compounds that inhibit MmpL3 without whole genome sequencing of isolated resistant mutants. Here we describe the identification of four new inhibitors that select for resistance mutations in *mmpL3.* Using these resistant mutants, we conducted a targeted whole-cell phenotypic screen of 163 novel Mtb growth inhibitors for differential growth inhibition of wild type Mtb as compared to a pool of twenty-four unique *mmpL3* mutants. The screen successfully identified six additional putative MmpL3 inhibitors. The compounds were bactericidal both *in vitro* and against intracellular Mtb. Mtb cells treated with these compounds were shown to accumulate trehalose monomycolate and have reduced levels of trehalose dimycolate, supporting MmpL3 as the target. The inhibitors were mycobacteria specific with several also showing activity against the non-tuberculosis mycobacterial species *M. abscessus.* Cluster analysis of cross resistance profiles generated by dose response experiments for each combination of 13 MmpL3 inhibitors against each of the 24 *mmpL3* mutants defined two clades of inhibitors and two clades of *mmpL3* mutants. Pairwise combination studies of the inhibitors revealed interactions that were specific to the clades identified in the cross-resistance profiling. Additionally, modeling of resistance substitutions to the MmpL3 crystal structure revealed clade specific localization of the residues to specific domains of MmpL3, with the clades showing differential resistance. Several compounds exhibited high solubility and stability in microsomes and low cytotoxicity in macrophages, supporting their further development. The combined study of multiple mutants and novel compounds provides new insights into structure-function interactions of MmpL3 and small molecule inhibitors.

## Introduction

In efforts to identify new tuberculosis (TB) antibiotics, whole cell-based phenotypic screens have been conducted against the pathogen *Mycobacterium tuberculosis* (Mtb). Over the last decade, several of these screens have identified MmpL3 as the proposed target for diverse small molecule inhibitors including AU1235, BM212, C215, DA-5, E11, indolecarboxamides, HC2091, NITD-349, PIPD1, Rimonabant, Spiro, TBL-140, THPP and SQ109^1, 2, 3, 4, 5, 6, 7, 8, 9, 10, 11, 12^. MmpL3 is an essential flippase responsible for transporting acetylated-trehalose monomycolate (TMM) synthesized in the cytoplasm to the pseudo-periplasmic space^13, 14, 15, 16, 17^. These TMMs are then converted into trehalose dimycolate (TDM) by the Ag85 complex in the cell envelope ^18^. MmpL3 is essential as evidenced by a pre-existing rescue allele being required to generate an *mmpL3* knockout^2, 14, 17, 19, 20, 21^, lack of mutants in high-throughput transposon mutagenesis screens^22, 23^, and studies that show rapid killing *in vitro* and *in vivo* in acute infection models when *mmpL3* expression is conditionally inhibited^14, 19^. This makes MmpL3 an attractive target for drug development, with one of its inhibitors, SQ109, currently in clinical trials ^24^.

MmpL3 inhibitors fall into diverse classes of chemical scaffolds ^25, 26, 27^, making it hard to computationally predict potential MmpL3 inhibitors based on chemical scaffolds. However, given the frequent finding of MmpL3 as a target, it is reasonable to expect that many new hits in a high throughput screen (HTS) may be acting against MmpL3. MmpL3 inhibitors have been identified by the isolation and sequencing of resistant mutants with single nucleotide variations (SNVs) mapping to the coding region of *mmpL3*, which is time-consuming and costly. Efforts to discover MmpL3 inhibitors using targeted approaches include generating hypomorphs, where a *mmpL3* knock down strain showed enhanced sensitivity to MmpL3 inhibitors, including AU1235 ^14^. However, this strain was also shown to be sensitive to isoniazid (INH) an inhibitor of InhA of the FAS-II pathway involved in mycolic acid synthesis, suggesting that while a *mmpL3* knockdown strain has robust screening potential for inhibitors of mycolic acid synthesis, maturation, and transport, such strains are not specific enough to identify inhibitors that selectively target MmpL3.

An alternative approach, employed in this study, is to use a pool of *mmpL3* resistant mutants to discover potential MmpL3 inhibitors. MmpL3 is a member of the resistance nodulation and division (RND) family of proteins, normally associated with efflux pumps in gram-negative bacteria ^2, 13, 17^. However, evidence suggests MmpL3 does not act as a general efflux pump in resistant backgrounds as resistant mutants do not differ in the amount of inhibitor isolated from cell fractions compared to WT Mtb ^2^. In further support that MmpL3 does not act as an efflux pump, the low level of cross resistance to compounds not associated with MmpL3 inhibition, including INH, suggests that MmpL3 does not act as a general efflux pump ^21^. This suggests that MmpL3 inhibitor resistant mutants could be used to screen for other potential MmpL3 inhibitors. The goal of this study was to discover MmpL3 inhibitors from a collection of 163 newly discovered, uncharacterized inhibitors of Mtb growth ^28^. Herein we describe the identification of four novel MmpL3 inhibitors by isolation of resistant Mtb mutants with mutations mapping to *mmpL3.* These twenty-four unique Mtb *mmpL3* mutant strains were then pooled into a single batch culture to conduct a targeted whole-cell phenotypic screen to identify six new scaffolds with reduced activity in the mixed mutant population as compared to the wild type. Cross resistance and compound interactions studies demonstrate specific structure function interactions between the molecules and MmpL3 and defined domains of MmpL3 associated with differential resistance to MmpL3 inhibitors.

## Results

### Identification of four new MmpL3 inhibitors by isolation of resistant mutants

Previously, two HTS were conducted, targeting the two component regulatory systems, DosRST and PhoPR^28, 29, 30^. In addition to inhibitors targeting these pathways, a series of compounds was identified that inhibited Mtb growth independent of the targeted pathways ^9, 28, 30^. A series of high throughput assays were then conducted to prioritize these compounds (Supplemental Figure 1) including confirming hits, testing for eukaryotic cytotoxicity in primary murine bone marrow-derived macrophages (BMMΦ, ≤10% cytotoxicity), and testing for the ability of the compounds to inhibit Mtb growth inside BMMΦ (≥25% growth inhibition). Results of these screens identified 216 compounds of which 163 commercially available compounds were purchased as fresh powders. In order to identify the mechanism of action of these Mtb growth inhibitors our lab selected several compounds with potent Mtb growth inhibition, both *in vitro* and in macrophages, as well as low eukaryotic cytotoxicity.

Four compounds of interest HC2060, HC2149, HC2169, and HC2184 (1-({1-[4-(Benzyloxy)-3-methoxybenzyl]piperidin-3-yl)carbonyl}azepane, N-[2-Methyl-6-(trifluoromethyl)pyridin-3-yl]-4-(trifluoromethyl)benzamide, Ethyl 3-{[(3,4-dihydro-2H-chromen-3-ylamino)carbonyl]amino}benzoate, and N-(2-Diethylaminoethyl)-N-(5,7-dimethyl-1,3-benzothiazol-2-yl)furan-2-carboxamide, respectively) (Figure 1a) had half maximal effective concentrations (EC_50_) ranging from 1.8 μM to 16.9 μM *in vitro* (Figure 1b, Table 1). All four compounds had bactericidal activity when measured at 20 μM (2x the initial screening concentration) (Figure 1c). To our knowledge, the structures of these compounds are unique from previously described inhibitors of Mtb growth.

**Figure 1:**
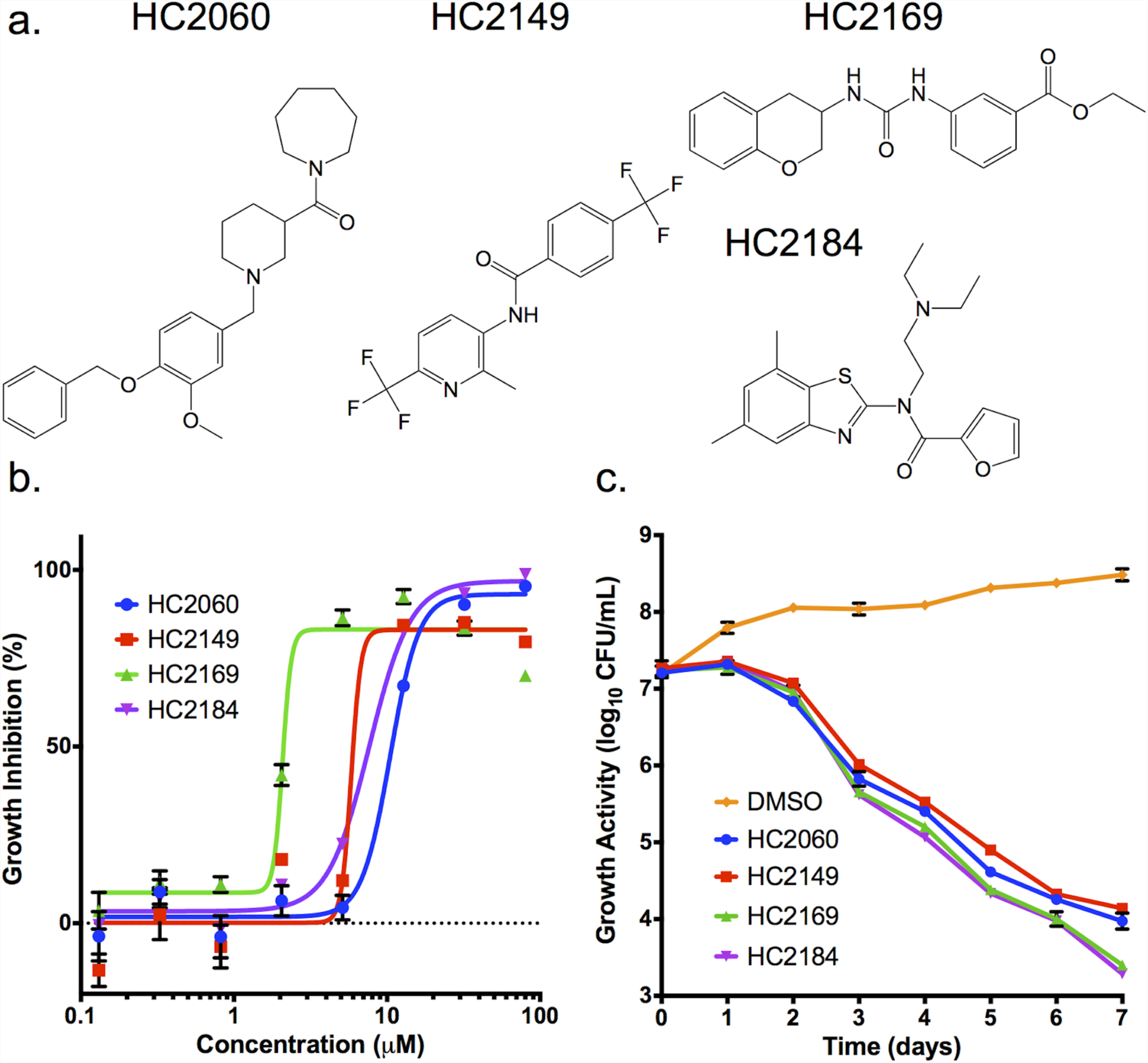
Four compounds inhibit Mtb growth in a dose and time dependent manner. **a)** Structures of HC2060, HC2149, HC2169 and HC2184. **b)** Inhibition of Mtb growth in a dose dependent manner. **c)** Killing of Mtb in a time dependent manner when treated at 20μM of the inhibitors. Error bars indicate the standard deviation from the mean. Experiments were conducted in biological triplicate.

**TABLE 1.** Characterization of MmpL3 Inhibitors

| Compound | WT EC<sub>50</sub><br>(μM) | *mmpL3* Mutant<br>Pool EC<sub>50</sub> (μM) | Relative<br>EC<sub>50</sub> | MΦ<br>EC<sub>50</sub> | Δψ<br>Disruption | Cytotoxicity<br>(CC<sub>50</sub>) (μM) | Solubility<br>(μM) | Microsome Stability<br>(% remain 30 min) |
| --- | --- | --- | --- | --- | --- | --- | --- | --- |
| HC2032 | 2.2 | >80 | >36 | 0.8 | Yes | >100 | 18 | 102 |
| HC2060 | 16.9 | >80 | >5 | 4.1 | No | >100 | >300 | 44 |
| HC2091 | 6.2 | >80 | >13 | 2.2 | No | >100 | >300 | 45 |
| HC2099 | 1.7 | 38.9 | 23 | <0.3 | No | >100 | 178 | 71 |
| HC2134 | 1.4 | >80 | >57 | 7.3 | Yes | >100 | 116 | N.D. |
| HC2138 | 4.0 | >80 | >20 | <0.3 | Yes | >100 | 66 | 122 |
| HC2149 | 6.6 | >80 | >12 | 3.6 | Yes | >100 | 131 | 138 |
| HC2169 | 1.8 | >80 | >44 | <0.3 | No | >100 | 17 | 168 |
| HC2178 | 3.8 | >80 | >24 | 2.0 | Yes | >100 | >200 | 4 |
| HC2183 | 3.2 | 59.9 | 19 | 3.0 | No | >100 | >200 | 25 |
| HC2184 | 7.6 | >80 | >11 | 0.7 | Yes | >100 | >300 | 30 |
| C215 | 11.2 | 57.5 | 5 | 4.0 | Yes | 14.3 | 87 | 62 |
| SQ109 | 2.4 | 6.9 | 2 | <0.3 | Yes | N.D. | N.D. | N.D. |

N.D. – Not determined
Relative EC<sub>50</sub> is fold difference between WT vs. *mmpL3* mutant pool.

To understand the mechanism of action of these four compounds, resistant mutants were isolated using solid agar plates (7H10 OADC) amended with 20 or 40 μM of each compound inoculated with 10^9^ CFU of Mtb (Erdman). Isolated mutants were tested for resistance via dose response curves. Confirmed resistant clones were isolated as single colonies and retested to confirm resistance (Supplemental Figure 2a-d). Genomic DNA was extracted from confirmed resistant mutant strains and the genomes were sequenced. Analysis of the genome sequences identified single nucleotide variants (SNVs) in all of the genomes in the coding region of *mmpL3* (Rv0206c, Supplemental Table 1). These mutations encoded for nonsynonymous mutations located throughout the gene (Supplemental Table 1, Supplemental Figure 2e). These findings suggest these compounds may be functioning as MmpL3 inhibitors.

### Modulation of TMM and TDM accumulation

MmpL3 is responsible for the transport of TMM across the inner membrane ^14, 15, 16, 18^. To determine if these compounds inhibited the activity of MmpL3, cultures of Mtb were grown in the presence of ^14^C-acetate and treated for 24 h with 20 μM of HC2060, HC2149, HC2169, HC2184, SQ109 or equal volumes of dimethylsulfoxide (DMSO). Radiolabeled lipids were isolated and analyzed by thin layer chromatography (TLC) (Figure 2a, Supplemental Figure 3a). The results of the lipid assay show that TMM accumulates in Mtb samples treated with the proposed MmpL3 inhibitors as well as the SQ109 treated samples. Additionally, TDM significantly decreased in cultures treated with HC2169 and HC2184 as well as the positive control SQ109 (Figure 2a and Supplemental Figure 3a). These results are consistent with previously described MmpL3 inhibitors ^1, 2, 3, 4, 6, 7, 8, 9, 26^ and support that these four compounds inhibit MmpL3 activity.

**Figure 2:**
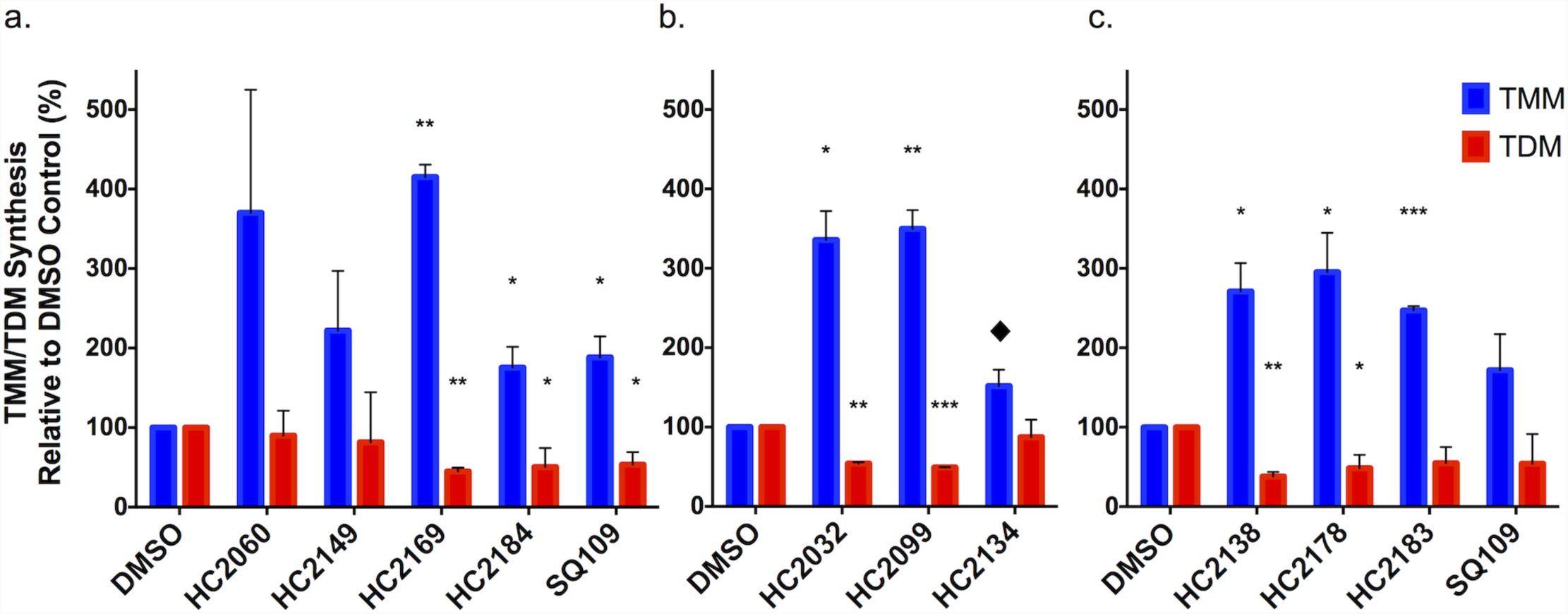
Modulation of TMM and TDM accumulation. **a)** Whole cell ^14^C-lipids from Mtb treated with 20μM of HC2060, HC2149, HC2169 and HC2184 show increased levels of TMM and decreased levels of TDM. **b and c)** Whole cell ^14^C-lipid from Mtb treated with 20μM of the six inhibitors identified by the targeted phenotypic screen show increased levels of TMM and decreased levels of TDM. Experiments were conducted in biological duplicate. In both experiments Mtb samples were treated with DMSO or 20μM SQ109. Error bars indicate the standard deviation. p-value < 0.05 (*), <0.005 (**), <0.001 (***). ♦ indicates values that just missed the cut off, HC2134 TMM p-value = 0.07. HC2060 and HC2149 missed significance cutoffs, but this may be due to the high variability in reps as the relative there was a >2 fold in difference for HC2060 and HC2149.

### Targeted whole cell phenotypic screen for MmpL3 Inhibitors

The identification of four new MmpL3 inhibitors, as well as the previously published inhibitor HC2091 (6), suggested additional MmpL3 inhibitors may exist in the prioritized 163-compound library of Mtb growth inhibitors (Supplemental Figure 1). Review of the known MmpL3 inhibitor scaffolds and those in our compound library identified HC2172 as the previously described MmpL3 inhibitor C215 ^7^. A recent study by McNeil *et al.*, showed that *mmpL3* mutant strains had low cross resistance against non-MmpL3 inhibitors^21^, suggesting that *mmpL3* mutants could be used to screen for MmpL3 inhibitors. Additionally, this study also showed that different mutations conferred varying levels of cross resistance between MmpL3 inhibitors. We therefore hypothesized that by pooling unique *mmpL3* mutant strains into a single mixed culture we could overcome limitations of cross resistance variability. For the targeted phenotypic screen, we directly compared percent growth inhibition (%GI) of either WT or a mixed *mmpL3* mutant pool consisting of twenty-four unique *mmpL3* mutant strains, including three strains previously described as resistant to HC2091 ^9^ (see Supplemental Table 2). The cultures were treated with 20μM of each of the 163-Mtb growth inhibitors as well as DMSO (negative control), Rifampin (RIF, positive control), Bedaquiline (BDQ), Clofazimine (CFZ), INH, para-aminosalicylic acid (PAS), H_2_O_2_, or SQ109 for a total of 171 different treatments (Supplemental Figure 4a and 4b). The results of this screen identified thirty-two compounds with 15% GI in the WT background and 1.5x reduced activity in the mixed *mmpL3* mutant background relative to the WT background (examples of positive hits are illustrated in red in Figure 3a). These hits were tested by dose response experiments conducted in both the WT and mixed *mmpL3* mutant background. In total, we identified thirteen compounds with reduced activity in the mixed *mmpL3* mutant background (Table 1, Supplemental Figure 5). Included in our confirmed hits were each of the five inhibitors used to generate the *mmpL3* mutant strains (HC2060, HC2091, HC2149, HC2169, and HC2184) and the two control compounds C215 and SQ109. The targeted screen also identified six novel inhibitors including HC2032, HC2099, HC2134, HC2138, HC2178, and HC2183 (ethyl 4-[(2E)-2-(4,7,7-trimethyl-3-oxo-2-bicyclo[2.2.1]heptanylidene)hydrazinyl]benzoate, 2-[(6-chloro-1H-benzimidazol-2-yl)sulfanyl]-N,N-di(propan-2-yl)acetamide, N-(2-methoxy-5-nitrophenyl)-1-oxo-4-phenylisochromene-3-carboxamide, 1-cyclohexyl-3-[4-[(2-fluorophenyl)methyl]-3-oxo-1,4-benzoxazin-7-yl]urea, 1-cyclooctyl-4-(2,5-dimethylphenyl)piperazine, and 2-[(6-methyl-1H-benzimidazol-2-yl)sulfanyl]-N,N-di(propan-2-yl)acetamide, respectively) (Figure 3b), which have not been previously described as MmpL3 inhibitors. The amount of resistance conferred by the mixed *mmpL3* mutant strains against each compound varied, with some compounds, like HC2032, HC2138, and HC2169 losing nearly all activity in the mutant background (Supplemental Figure 5) as indicated by the high Relative EC_50_ (fold difference between *mmpL3* mutant pool and WT) values of >36, >20, and >44 (Table 1). Despite the high activity of SQ109 in the WT background, the relative EC_50_ was only 2 (Table 1); however, this observation is consistent with previous studies which only report marginal increases in MIC values in *mmpL3* mutant backgrounds ^8, 9, 21, 26^. Included in our hits were two urea-based compounds HC2138 and HC2169 (Figure 1 and Figure 3b). These urea-based compounds have structures reminiscent of the adamantyl-urea MmpL3 inhibitor AU1235 ^2^. Additionally, two of the compounds identified in the screen HC2099 and HC2183 had high structure similarity.

**Figure 3:**
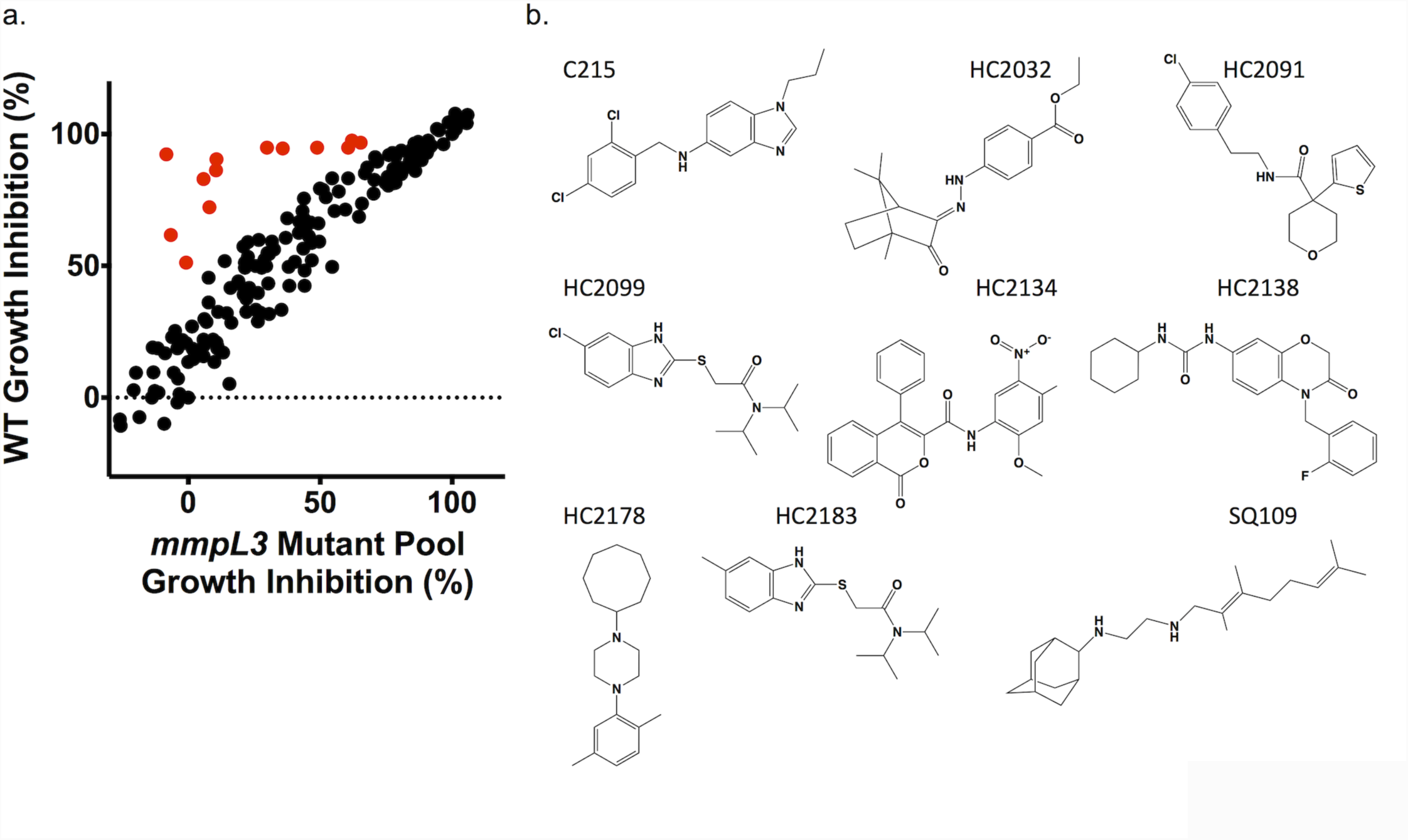
A targeted whole cell phenotypic screen identifies six new MmpL3 inhibitors. **a)** Results of a direct head to head comparison of percent growth inhibition of WT Mtb or a pooled *mmpL3* mutant population treated with 20μM of 163 compounds. Additional treatments included 0.5μM of BDQ, CFZ, INH, PAS, SQ109 or 0.03% H_2_O_2_. Examples of hit compounds with reduced activity in the MmpL3 mutant pool are shown in red. **b)** Structures of the confirmed hits from the screen, including six new compounds HC2032, HC2099, HC2134, HC2138, HC2178, and HC2183. Previously described compounds include C215, HC2091 and SQ109. Radio-TLCs are shown in Supplemental Figure 3.

The compounds were also tested for eukaryotic cytotoxicity, solubility and stability in mouse microsomes, and the structures were confirmed by mass spectrometry (Table 1). The compounds exhibited low cytotoxicity (>100µM), consistent with our secondary assay screening. Compounds exhibited varying levels of solubility with HC2169 and HC2138 showing lower solubility (66μM and 17μM respectively) but high microsome stability (122% and 168% respectively), and compounds like HC2183 showing high solubility (>200μM) but low microsome stability (25%). Interestingly HC2099, which has high structure similarity to HC2183 showed higher solubility (178μM) and higher microsome stability (71%). Several of the compounds (e.g. HC2091, HC2099, HC2138 and HC2149), exhibited favorable solubility and microsome stability, with no observed macrophage cytotoxicity, supporting their potential for further development.

The phenotypic screen was selective as it did not identify any of the control treatments known to not target MmpL3 including BDQ, INH, PAS, H_2_O_2_, or HC2051, a proposed Pks13 inhibitor (given its similarity to the TAM16 ^31, 32^). To confirm the specificity of our screen, we conducted dose response studies in both the WT and mixed *mmpL3* mutant background for each of the aforementioned inhibitors, as well as RIF. Results of the dose response studies did not identify any significant levels of resistance to these compounds in the mixed *mmpL3* mutant background (Supplemental Table 3, Supplemental Figure 6). This was true for both inhibitors of mycolic acid synthesis and maturation (INH and HC2051), suggesting our screen was specific for inhibitors of MmpL3. Consistent with previous results, we identified increased susceptibility to RIF treatment in the mixed *mmpL3* mutant background^21^ (Supplemental Table 3, Supplemental Figure 6). The dose response profiles for BDQ, CFZ, and PAS did not show any differences in susceptibility, further supporting that *mmpL3* mutations do not confer resistance through general efflux.

### Modulation of TDM, membrane potential and viability

To determine if the six compounds identified in the screen can inhibit MmpL3 activity, we examined accumulation of TMM and TDM as described above. The inhibitors modulated mycolic acid accumulation in whole cell extracts, with lipids for all treatments showing a significant accumulation in TMM (except for HC2134) and treatment with HC2032, HC2099, HC2138, and HC2178 showing a significant decrease in TDM relative to the DMSO control samples (Figure 2b and 2c, Supplemental Figure 3b). A recent report has shown that because MmpL3 activity is dependent on the proton motive force (PMF), disruptors of PMF, such as the protonophore carbonyl cyanide m-chlorophenyl hydrazine (CCCP) can also modulate MmpL3 activity ^26^. Studies have suggested that some proposed MmpL3 inhibitors such as SQ109 and E11 may indirectly target MmpL3 through disruption of the membrane potential ^6, 15, 26^. To determine if the newly identified inhibitors disrupt membrane potential (ΔΨ) we conducted dose response studies using a DiOC_2_-based assay. Some compounds, including HC2060, HC2169, and HC2183 did not disrupt membrane potential (Table 1, Supplemental Figure 7), while others, such as HC2032, HC2099, HC2134, HC2138, HC2149, HC2178, HC2184 and C215 did disrupt membrane potential (Table 1, Supplemental Figure 7). Consistent with previous observations HC2091 did not disrupt membrane potential, while SQ109 did disrupt membrane potential (Table 1, Supplemental Figure 7) ^9, 15, 26^. Surprisingly, there were differences in outcome for the two urea containing compounds HC2169 and HC2138 as well as between HC2099 and HC2183 which only differ by a chloro and methyl substitution, respectively. The results for HC2138 and HC2169 is also interesting because the previously described urea-based MmpL3 inhibitor, AU1235, does not disrupt the membrane potential^15, 26^. These results suggest that the ability to disrupt membrane potential is highly structure specific.

Because MmpL3 is essential for viability of replicating bacteria, we tested these compounds for bactericidal activity using a firefly luciferase (*luc*) reporter strain of Mtb in conjunction with a luciferase assay. This assay relies on active luciferase generated by the reporter Mtb strain, and the presence of ATP which is generated in living cells, but rapidly hydrolyzed in lysed cells. All of these compounds showed bactericidal activity (Supplemental Figure 8). These results suggest that the growth inhibition is due to compounds killing Mtb in a dose dependent manner. The bactericidal activity of these inhibitors is consistent with these compounds targeting MmpL3 which is essential for cell viability ^14, 19^.

### Spectrum of activity

While MmpL3 is conserved in mycobacteria, functional orthologs are not found in other bacteria and fungi. Despite this, several proposed MmpL3 inhibitors including BM212, THPP, and SQ109 have been shown to inhibit multiple bacterial and fungal species ^8, 33, 34, 35^ while other MmpL3 inhibitors including HC2091, AU1235, and indolecarboxamides are specific to mycobacteria. To define the spectrum of activity, the compounds were tested against several diverse species including *Staphylococcus aureus, E. coli, Pseudomonas aeruginosa, Proteus vulgaris*, and *Enterococcus faecalis* (Table 2). For HC2032, HC2060, HC2099, HC2149, HC2169, HC2178 and HC2184, even at high concentrations (200μM), no inhibition was observed against non-mycobacteria. However, these inhibitors were positive for activity against other mycobacteria, including the pathogenic non-tuberculosis species *M. abscessus* and the saprophytic species *M. smegmatis* (Table 2). For example, HC2091, HC2099, and HC2134 exhibited MIC_50_ of 6.25 µM, 25 µM and 12.5 µM against *M. abscessus*, respectively. Additionally, all of the MmpL3 inhibitors tested, except for HC2149, were active against *M. smegmatis*. This suggests that most of the inhibitors are specific for mycobacteria and may be effective against diverse mycobacterial species. The observation that HC2134 and C215 are active against non-mycobacterial species has been observed with other MmpL3 inhibitors ^24, 34, 35^ and may be due to a non-specific activities such as PMF disruption.

**TABLE 2.** EC_50_ Values for Spectrum of Activity of MmpL3 Inhibitors*

|  | HC2032 | HC2060 | HC2091 | HC2099 | HC2134 | HC2138 | HC2149 | HC2169 | HC2178 | HC2184 | C215 |
| --- | --- | --- | --- | --- | --- | --- | --- | --- | --- | --- | --- |
| Mtb – Erdman | 3.0 | 14.8 | 7.0 | 5.3 | 2.1 | 2.3 | 11 | 2.4 | 3.7 | 8.9 | 16.2 |
| Mtb – CDC1551 | 2.4 | 12.8 | 6.3 | 4.8 | 1.5 | 2.0 | 10.5 | 2.2 | 2.3 | 7.6 | 14.3 |
| Mab | 34.5<sup>a</sup> | 13.6<sup>a</sup> | 96.5<sup>a</sup> | 81.8<sup>a</sup> | 81.9<sup>a</sup> | 8.2<sup>a</sup> | -28<sup>a</sup> | -36<sup>a</sup> | 13.5<sup>a</sup> | 7.0<sup>a</sup> | 2.3<sup>a</sup> |
| Msm | 2.2 | 80 | 20<sup>9</sup> | 0.9 | 1.8 | N.D. | >200 | 13.2 | 4.5 | 85.7 | >200 |
| *S. aureus* (1) | >200 | >200 | >200<sup>9</sup> | >200 | 51.0 | N.D. | >200 | >200 | >200 | >200 | 15.1 |
| *S. aureus* (2) | >200 | >200 | >200<sup>9</sup> | >200 | 51.8 | N.D. | >200 | >200 | >200 | >200 | 22.8 |
| *E. coli* | >200 | >200 | >200<sup>9</sup> | >200 | >200 | N.D. | >200 | >200 | >200 | >200 | >200 |
| *P. vulgaris* | >200 | >200 | >200<sup>9</sup> | >200 | 35.1 | N.D. | >200 | >200 | >100 | >200 | >200 |
| *E. faecalis* | >200 | >200 | >200<sup>9</sup> | >200 | >200 | N.D. | >200 | >200 | >200 | >200 | 34 |
| *P. aeruginosa* | >200 | >200 | >200<sup>9</sup> | >200 | >200 | N.D. | >200 | >200 | >200 | >200 | >200 |

\*All values are EC<sub>50</sub> except for *M. abscessus* data which are single concentration % inhibition data
a – Growth Inhibition (%) at 20μM
9 – Reference number of previously published data
Mab, *M. abscessus*; Msm, *M. smegmatis*
*S. aureus* (1) – *S. aureus* ATCC29213, *S. aureus* (2) – *S. aureus* ATCC25923
N.D. – not determined

### Activity against intracellular Mtb

The compounds were tested against Mtb growing in BMMΦ using a luciferase expressing Mtb strain. BMMΦ were infected with Mtb and treated with the inhibitors for six days across a range of concentrations (200 – 0.3µM). The BMMΦ EC_50_ values are summarized in Table 1 and Supplemental Figure 9. The results of the assay show that many of the inhibitors have bactericidal activity in MΦ several magnitudes lower than the eukaryotic cytotoxicity CC_50_, supporting a high selectivity index. The identification of bactericidal effects against Mtb in BMMΦ is consistent with genetic knockdown studies that show *mmpL3* is essential for actively replicating bacteria (4,5).

### Cross resistance profiles indicate specific MmpL3 protein-inhibitor interactions

While the results of the screen showed potential for rapid identification of MmpL3 inhibitors, the screen relied on the use of a mixed mutant population. To resolve this issue, we conducted dose response studies for each combination of the twenty-four unique *mmpL3* mutants against each MmpL3 inhibitor identified from the screen (with WT Mtb as a control). Because there was a complete lack of activity for compounds like HC2169 against HC2169-specific resistant mutants (Supplemental Figure 2c), units of measure such as EC_50_ and MIC cannot be calculated, or are not a good measure for comparing responses. Instead, we used the area under the curve (AUC) for each dose response in the *mmpL3* mutant backgrounds relative to the AUC for the WT response for a given treatment (Supplemental Table 4). Because the compounds have differences in potency, the AUC for the WT for each treatment differs and to account for this issue we normalized our values by Z-score for each treatment^36^. Cluster analysis grouped the data based on both treatment effectiveness and resistance conferred by each *mmpL3* mutant. The resulting cluster-gram (Figure 4) shows that both compounds and *mmpL3* mutant strains, denoted by the amino acid substitutions, fall into distinct clades. The compounds fall into two distinct clades, Clade A (Red), which contains HC2134, HC2138, HC2149, HC2169, and Clade B (Green) which contains HC2032, HC2060, HC2091, HC2099, HC2178, HC2183, HC2184, C215, and SQ109. The identification of two distinct clades of compounds, suggested that the compounds may be interacting with the MmpL3 protein in distinct ways.

**Figure 4:**
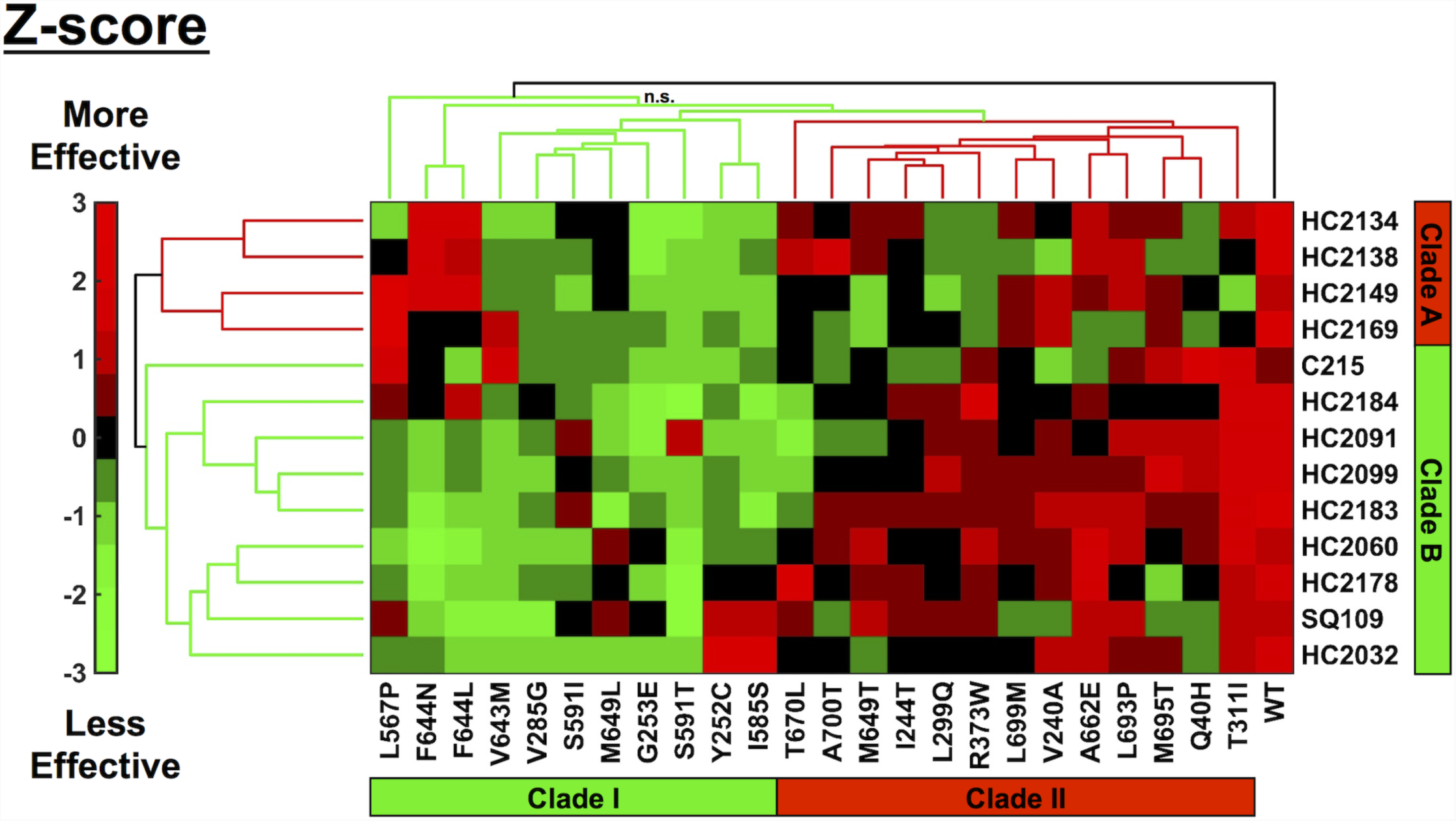
Cross resistance profiling identifies clustering of compounds and mutations. Cluster analysis of cross-resistance profiling of twenty-four *mmpL3* strains treated with each of the thirteen MmpL3 inhibitor normalized by Z-scoring by treatment. Compounds clustered into two clades: Clade A and Clade B. Mutant strains, denoted by amino acid substitution, clustered into two clades: Clade I and Clade II. Colors are based on Z-score normalization of treatment, green indicates when treatments were less effective and red indicates when treatments were more effective than the average (black). n.s. indicates a branch where the approximate unbiased (AU) value was < 75. All other branches were significant based on bootstrap AU values > 75.

The resistance mutations also showed specific clustering. Cluster analysis of the strains showed that WT clustered on its own and the mutants formed a large complex cluster. Within this large cluster, the *mmpL3* mutant strains formed into two sub-clades; Clade I (Green) which conferred relatively high resistance (lower inhibitor effectiveness) and Clade II (Red) which conferred relatively low resistance (higher inhibitor effectiveness). Clade I, which contained eleven *mmpL3* mutant strains denoted as Y252C, V285G, G253E, L567P, I585S, S591I, S591T, V643M, F644N, F644L, and M649L. Clade II consisted of the remaining thirteen *mmpL3* mutant strains denoted as Q40H, V240A, I244T, L299Q, T311I, R373W, M649T, A662E, T670L, L693P, M695T, L699M, and A700T. Surprisingly M649T fell into Clade II mutations, this was striking as the *mmpL3* mutant denoted as M649L was clustered with the Clade I *mmpL3* strains.

### Pairwise Combination Studies using DiaMOND

We hypothesized that the clustering of compounds into two clades was due to their having distinct interactions with MmpL3; therefore, combination treatments may reveal antagonistic, additive or synergistic interactions. In order to test this hypothesis in whole cell Mtb, we used the recently described <u>dia</u>gonal <u>m</u>easurement of *<u>n</u>*-way <u>d</u>rug interactions (DiaMOND) approach ^37^. RIF was included as a control for these assays, as this drug has been shown to be synergistic when tested with other MmpL3 inhibitors such as AU1235 and SQ109 ^38, 39^. The results of DiaMOND, shown in Figure 5, identified synergistic interactions (FIC_2_ < 1.0) between all combinations of MmpL3 inhibitors and RIF. Additionally, the results identified mostly additive interactions (FIC_2_ = 1.0) consistent with the compounds sharing a single target. Interestingly, all combinations between MmpL3 inhibitors and the compounds HC2134, HC2138, HC2149, and HC2169 showed antagonistic interactions (FIC_2_ > 1.0). These four compounds were clustered together in Clade A in the cross-resistance profiles described above (Figure 4). This antagonistic relationship further supports that the Clade A compounds are distinct from the Clade B compounds. Another observation from the DiaMOND assay is that pairwise combinations of compounds HC2060, C215, and SQ109 all had synergistic interactions (Figure 5). The reason for this observation is not clear as the compounds did not have differential cross resistance profiles (Figure 4). Interestingly, combinations of HC2060 and C215, but not SQ109, with the Clade A compounds HC2139 and HC2169 did not reveal antagonist interactions, but instead additive interactions (Figure 5). This finding supports that these HC2060 and C215 compounds interact with MmpL3 in a manner distinct from the other compounds.

**Figure 5:**
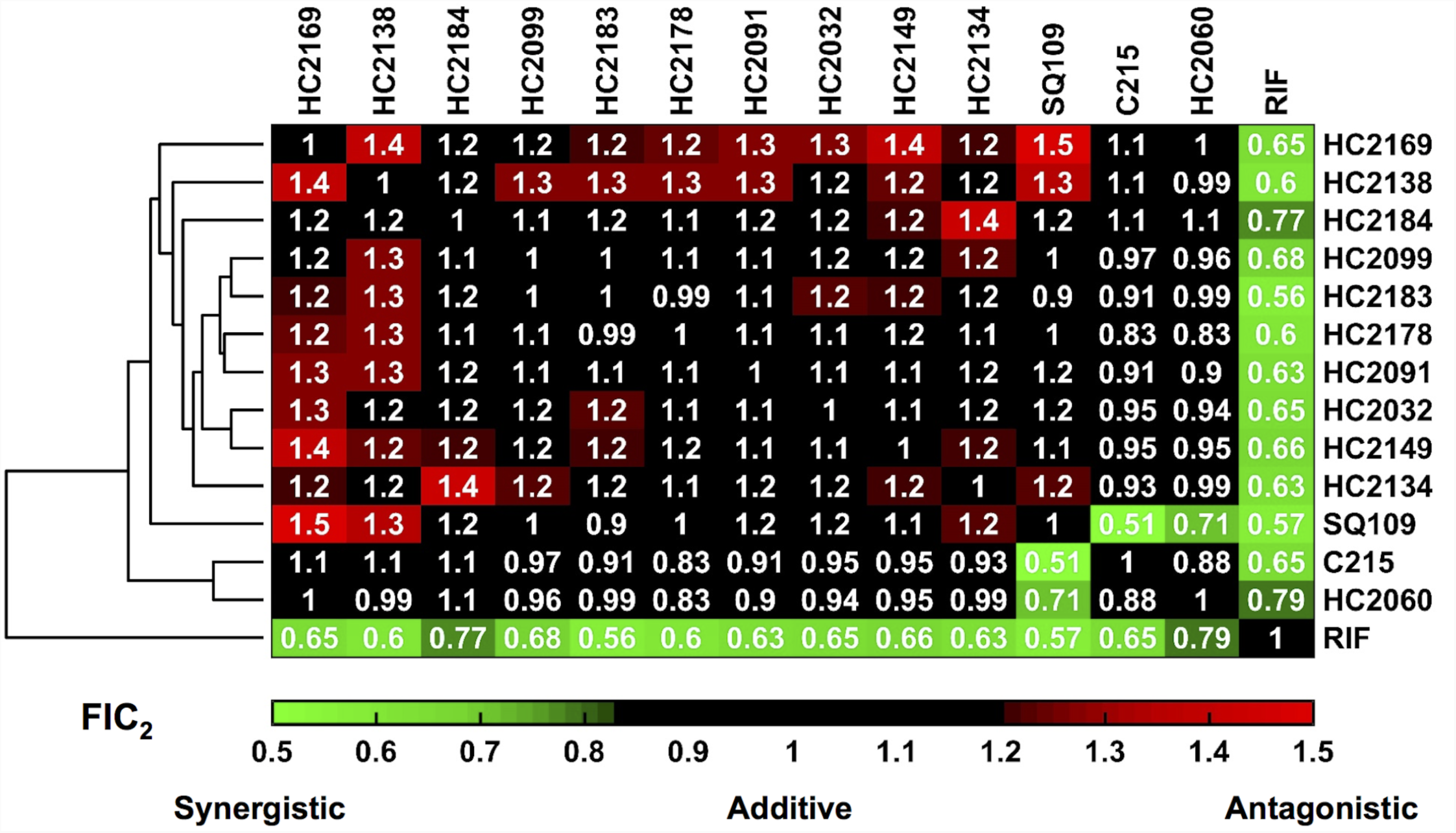
DiaMOND Analysis Identifies Additive, Synergistic and Antagonist Inhibitor Interactions. Hierarchical cluster analysis of DiaMOND-based pairwise inhibitor interactions of all combinations of MmpL3 inhibitors and RIF identifies additive (FIC_2_ 0.82-1.18) antagonistic (FIC_2_ > 1.18) and synergistic (FIC_2_ < 0.82) interactions.

## Discussion

The cross-resistance profiles showed that the *mmpL3* mutant strains clustered separately into two clades, Clade I and Clade II, with WT clustering on its own as an outgroup (Figure 4). Recently, the crystal structure of *M. smegmatis* MmpL3 has been solved by two independent groups^12, 40^. In order to understand this observation, we generated a 3D model of the Mtb MmpL3 protein aligned to the *M. smegmatis* MmpL3 structure^41^ (C-score 0.17, RMSD 8.4 ±4.5Å). Substitutions from the *mmpL3* mutant strains used in the cross-resistance profiles are highlighted in the model (red, green, and blue) (Figure 6). Consistent with previously described resistant strains of Mtb, the majority of the substitutions, localized along the central vestibule with the exception of T670, R373, and A662 which did not align along the central vestibule of the model (Figure 6)^17^. This vestibule is conserved amongst the RND family of proteins and is responsible for the proton translocation that drives protein activity ^17, 26^. To understand the clustering pattern of the cross-resistance profiling we highlighted the mutations based on their clade, revealing that the two distinct clades separated spatially in the model. The substitutions of Clade I (Green), that conferred higher resistance, localized towards the cytoplasmic face of the protein. While substitutions of Clade II (Red), which generated lower resistance, localized into two separate locations: i) towards the pseudo-periplasmic face of the protein; and, ii) another region which does not line the central vestibule (Figure 6). Structure function profiling by Belardinelli and colleagues ^17^ had previously described seven essential residues for MmpL3 function (D251, S288, G543, D640, Y641, D710, and R715) that clustered in a single domain ^17^. This study also modeled substitutions commonly identified from resistant mutants to multiple inhibitors to this same region. To determine if the two clades separated based on their approximation to this essential region, we highlighted these seven residues in the model (Supplemental Figure 10). Notably, the two clades separate based on their proximity to these residues, with Clade I substitutions localizing in the same region as the essential residues and Clade II substitutions localizing distally from the essential residues. This finding suggests that the strength of resistance conferred by a mutation is dependent on the proximity of the substitution to residues essential for MmpL3 function.

**Figure 6:**
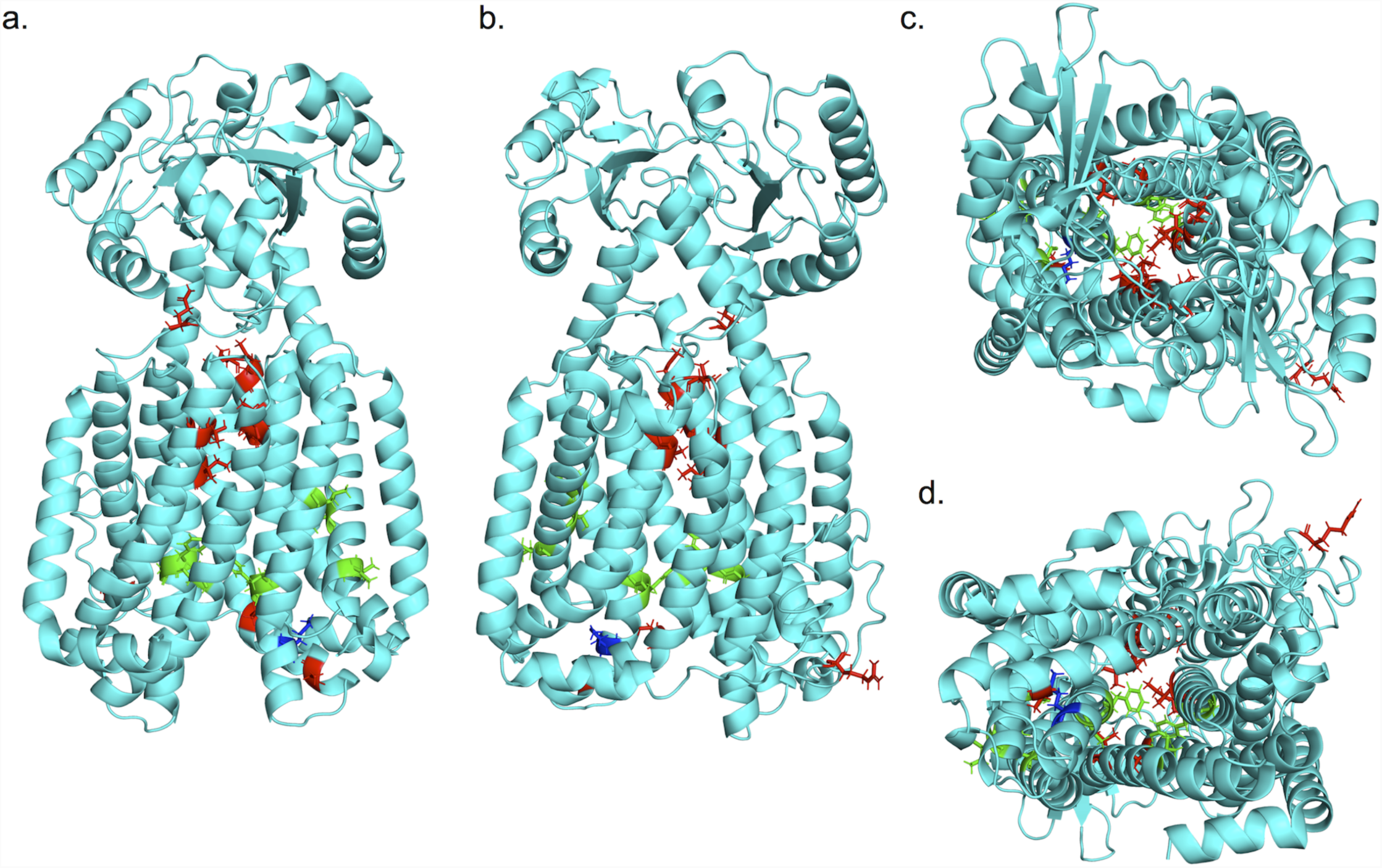
Mutation substitutions cluster according to cross resistance clades. **a-d)** Front, back, top, and bottom (respectively) views of an I-TASSER predicted structure of Mtb MmpL3 based on *M. smegmatis* MmpL3 structure (PDB: 6AJH). Substitutions conferred by mutations in *mmpL3*. Substitutions are colored based on clade from cross resistance profiling, Clade I substitutions (green), Clade II substitutions (red), or M649 (blue) which fell into both clades depending on substitution. The model shows a truncated version (732/944aa) of the MmpL3 protein lacking the C-terminal tail.

Genome sequences of the isolated resistant mutants identified a total of 21 unique mutations in *mmpL3*. These mutations translated to substitutions that were a mix of ones previously described and novel to this study. Included in this list were substitutions that had previously been described including G253E, Y252C, L567P, S591I, V643M, F644L, L699M ^1, 2, 5, 6, 8, 9, 21, 31^. Mutations that were unique to this study included ones in positions in V240, I244, V285, L299, T311, R373, I585, A662, and L693. We also isolated mutations that had previously been described to occur in positions Q40, Y252, G253, L567, S591, F644, M649, and L699 ^1, 2, 5, 6, 8, 9, 21, 31^. However, the exact substitutions in several of these strains’ positions including Q40H, S591T, F644N, and M649T were unique to this study. Our cross-resistance profiling found that G253E, V285G, S591I, S591T, L699M, and A700T, conferred pan resistance including against SQ109. That the number of compounds proposed to target MmpL3 and the large number of substitutions that confer resistance highlights the importance of identifying combinations of drugs that would reduce the frequency of resistance.

The favorable properties of many of these compounds, including low cytotoxicity, high solubility and microsome stability, and activity in macrophages, suggests that these compounds warrant further development as new therapeutics. It is also possible that combinations of these scaffolds may be developed in a single molecule that can function to reduce the frequency of resistance. Three of the compounds used to isolate resistant mutants in this study, HC2149, HC2169 and HC2184, had a frequency of resistance (FoR) of 3 x 10^-7^, which is similar to the FoR of other MmpL3 inhibitors that have a FoR ranging from 10^-7^ to 10^-8^ ^2, 3, 4, 5, 6, 21^. That the FoR for HC2184 was the same as the FoR for HC2149 and HC2169 is interesting as the cross-resistance profiles and results of DiaMOND analysis suggested that these compounds interact differently with MmpL3. While the antagonistic interactions identified by DiaMOND suggest that scaffold combinations may lower the activity of a single inhibitor, antagonistic drug combinations have been proposed to decrease the rate of resistance ^42, 43^. It therefore may be possible to design a single inhibitor that fuses more than one scaffold to decrease the rate of resistance. This hypothesis could initially be tested by conducting pairwise combination studies examining for synergistic reductions in the FoR. Given the relative ease of resistance occurring to MmpL3 inhibitors, a reduced FoR could be a valuable new property for this class of inhibitors.

Over the past decade, many MmpL3 inhibitor of various chemical scaffolds have been described. The proposed target of these inhibitors has been driven by the mapping of resistance mutations to *mmpL3*. The screening platform we describe here greatly accelerates target identification of such inhibitors. The use of a diverse pool of unique *mmpL3* mutants, rapidly identified inhibitors of MmpL3 activity, as demonstrated by their ability to modulate TDM and TMM accumulation. A subset of these inhibitors was shown to disrupt membrane potential, and potentially the PMF which energizes MmpL3 activity. Two recent studies have suggested that two MmpL3 inhibitors, SQ109 and E11, indirectly inhibit MmpL3 by targeting the PMF despite the isolation of *mmpL3* resistant mutants ^6, 15^ and co-crystallization of SQ109 to MmpL3^12^. It is therefore possible that some of these new compounds inhibit MmpL3 indirectly by dissipation of the PMF. Notably, the narrow spectrum of activity of most of the isolated compounds for mycobacteria supports that the target is mycobacterium specific, and not a general target that when bound dissipates PMF.

## Methods

### Media and Growth Conditions

Unless otherwise specified, streptomycin resistant strains of Mtb Erdman or CDC1551 were cultured in 7H9 media supplemented with 10% OADC (v/v) with 0.05% Tween-80 (v/v) in standing T25, T75 or T150 flasks at 37 °C with 5% CO_2_. Spectrum of activity studies in different bacterial species (Table 2) were conducted as described by Coulson et al., ^30^, with the exception of the *M. abscessus* studies which are described in the supplemental methods.

### Dose Response Curves

Mtb was grown in rich medium to an OD_600_ of 0.5-1.0. Cultures were diluted to an OD_600_ of 0.1 in 7H9 medium and aliquoted into black walled clear bottom 96 well assay plates. Compounds were tested between 80-0.13 μM with 2.5-fold dilutions, controls included DMSO and 3 μM RIF. Plates were placed in zip lock bags with moistened paper towels and incubated at 37°C for 6 days. Plates were read on a PerkinElmer Enspire plate reader. %GI was calculated using DMSO and RIF as 0% and 100% inhibition, respectively. Dose responses were conducted in biological triplicate and repeated at least once. Significant differences of EC_50_ were compared using 95% confidence intervals.

To examine the spectrum of activity of the MmpL3 inhibitors, the EC_50_ of each compound was also determined for *M. smegmatis* and other nonmycobacteria, including *S. aureus, E. coli, P. aeruginosa, P. vulgaris*, and *E. faecalis*. Tests were performed in 96-well plates in LB broth with shaking at 37°C, with the exception of *E. faecalis*, which was grown in brain heart infusion medium in standing flasks at 37°C, and *M. smegmatis*, which was also grown standing at 37°C in LB broth with 0.05% Tween-80. Culture was diluted to a starting OD_600_ of 0.05. Bacteria were incubated in the presence of an 8-point (2-fold) dilution series of each inhibitor ranging from 200 μM to 1.5 μM for 6 h, except for *M. smegmatis*, which was incubated for 72 h. Growth was monitored by measuring optical density and normalized based on kanamycin (100% growth inhibition) and DMSO (0% growth inhibition) controls, with the exception of *P. aeruginosa*, for which 10 μg/mL tobramycin was used as the control for 100% growth inhibition. The experiments were performed with three technical replicates per plate. EC_50_s were calculated based on a variable-slope four-parameter nonlinear least-squares regression model in the GraphPad Prism software package (version 8).

### Kinetic Kill Curves

Mtb was cultured in 7H9 medium to an OD_600_ of 0.5-1.0 and diluted to an OD_600_ of 0.1. In triplicate, diluted samples were aliquoted into 96 well plates and inoculated with 20 μM concentrations of each compound with DMSO as a negative control. Plates were placed in zip lock bags with moistened paper towels and incubated at 37 °C. Daily samples were taken and serial diluted in 96 well plates using 1x PBS + 0.05% Tween-80 (v/v) and plated on 7H10 quadrant plates supplemented with OADC (10% v/v). Plates were incubated at 37°C and colonies were counted to calculate CFU/mL. Experiments were conducted in biological triplicate and repeated at least twice.

### Isolation of Resistant Mutants

Mtb was grown to an OD_600_ of 0.6-1.0 and samples were resuspended in fresh media for a final cell count of 2 x 10^9^ cells/ml. Cell pellets were resuspended in 0.5 ml of 7H9 medium and plated on 7H10 OADC plates supplemented with 20 μM or 40 μM concentrations of HC2060, HC2149, HC2169, and HC2184. Plates were incubated at 37 °C until isolated colonies appeared. Colonies were picked and inoculated into 5 ml of 7H9 medium in T25 standing flasks and grown to an OD_600_ 0.5-1.0. Samples were taken and tested for resistance using dose response curves described above along with WT grown to an OD_600_ of 0.6-1.0 and 3 μM RIF and DMSO were used as controls. Samples were also serial diluted as described above and plated for colony purified single colony isolates on X-plates containing 7H10 OADC. Single colony isolates were picked and inoculated into 5 ml of 7H9 OADC in T25 flasks. Resistance was re-confirmed using the same methods described above. Differences in EC_50_ values were deemed significant based on the 95% confidence intervals.

### Whole Genome Sequencing and Analysis

Whole genome sequencing was performed as previously described^44^. Briefly cultures of single colony isolates were grown to an OD_600_ ~1.0 and pelleted. Genomic DNA was extracted and sequenced by Illumina-based whole genome sequencing at 250 bp reads. Sequencing results were analyzed using the GATK workflow for the identification of single nucleotide variations^45^.

### TMM and TDM accumulation assay

The lipid assay was carried out as previously described(6). Briefly, 30 ml cultures of Mtb was cultured to an OD_600_ of 0.6. Samples were diluted to an OD_600_ of 0.1 in 8 ml cultures in T25 flasks. Cultures were inoculated with 8 μCi of ^14^C-acetate. Cultures were co-inoculated with 20 μM samples of MmpL3 inhibitors and then incubated for 24 hours before performing lipid extraction as previously described^9^. Total extractable lipid ^14^C-incorporation was determined by scintillation counting, and 5,000 cpm of lipids were separated on TLC plates with a 24:1:0.5 Chloroform:Methanol:H_2_O solvent system. TLCs plates were imaged using a Typhoon FLA 7000 and images were quantified using IQ image quantifying software. Experiments were conducted in biological duplicate. Comparison to the DMSO controls was conducted using the T-test.

### Targeted whole cell phenotypic screening

Each *mmpL3* mutant was cultured independently in 8 ml of 7H9 medium in T25 standing flasks to an OD_600_ of 0.6 – 1.0. Mutant cultures were separately back diluted to an OD_600_ of 0.6 in 1.5 ml of 7H9 medium in 2 ml screw cap tubes. The contents of each tube were mixed into a single batch culture in a T75 culture flask. The mixed mutant culture was allowed to recover overnight (~8 hours) at 37°C. Samples of Mtb Erdman (WT, OD_600_ = 0.6) and the mixed mutant population were back diluted to an OD_600_ of 0.1 in 7H9 medium. WT and mutant pools were aliquoted, in technical duplicate, into separate clear bottom black walled 96 well plates. Samples of WT and mixed mutant cultures were inoculated with 20 μM of each of the 163 compounds from the small molecule library. Additional treatments included 0.5 μM samples of *para*-amino salicylic acid (PAS), SQ109, bedaquiline (BDQ), isoniazid (INH), clofazimine (CFZ), as well as DMSO and 0.3 μM RIF. Percent growth inhibition (%GI) of WT and mutant mix population were calculated for each treatment and hits were defined as 1) compounds with at least 15% GI in the WT background and 2) 1.5 fold decreased inhibition in the mutant pool relative to the WT background. The hit compounds were confirmed by conducting dose responses curves of screen hits as described above against WT and *mmpL3* mutant pools. Dose response curves were conducted in technical duplicate and differences between the WT and *mmpL3* mutant pool was deemed significant based on 95% confidence interval. Confirmed hits were reassessed with similar results.

Cross resistance studies were conducted by generating dose response curves for every combination of MmpL3 inhibitor and each *mmpL3* mutant, and WT Mtb strain, CDC1551 or Erdman depending on the background of the *mmpL3* strain (for a total of 338 dose response curves). Cross resistance dose responses were conducted singly, unless the dose response identified increased sensitivity in the *mmpL3* mutant background, in which case the responses were re-examined using dose responses carried out in biological duplicate. The dose responses were then used to calculate the area under the curve (AUC) using Prism 8 software using the default setting. AUCs were compare to the respective WT strains by dividing the AUC of the *mmpL3* strain by the respective WT parent strain. AUC fractions were then standardized by treatment by Z-scoring ^19^. Z-score standardized data was then clustered in MatLab by hierarchical agglomerative clustering using the *clustergram* function with default settings (Euclidean distance model, Average linkage clustering). Hierarchical agglomerative clustering using bootstrapped data was conducted in R using *pvclust* (nboot = 1000) using the Euclidean distance model and average linkage clustering ^46^.

### Membrane Potential Assays

The DiOC_2_ membrane potential assay was carried out as previously described ^1, 9^. Briefly, Mtb Erdman cells were labeled with 30 μM DiOC2 (Thermo Scientific) in 1 ml of 1X phosphate-buffered saline (PBS) (pH 7.4), supplemented with 50 mM KCl, and incubated at 37°C for 15 min. Cells were washed twice and suspended in 1X PBS at a final concentration of an OD_600_ of 0.2, and 200 μL of labeled cells were aliquoted to 96-well plates and treated with each of the MmpL3 inhibitors at 80 μM, 20 μM, or 5 μM concentrations. Samples were also treated with DMSO (negative control) or 25 μM CCCP (Sigma-Aldrich) (positive control). Each treatment included three technical replicates per plate. The kinetics of fluorescence (excitation, 485 nm; emission, 610 nm/515 nm) was measured every 2 min for 60 min. The red/green (610 nm/515 nm) fluorescence intensity ratio was calculated and used to quantify membrane potential. The experiment was repeated at least twice with similar results. The error bars represent the standard deviation of the geometric mean.

### Bactericidal Activity *in vitro* and in macrophages

An Mtb CDC1551 strain with a chromosomally encoded firefly luciferase ^47^ was grown to an OD_600_ of 0.6 – 1.0 in rich medium. For *in vitro* experiments cultures were diluted to an OD_600_ of 0.1 and aliquoted at 100μL in white walled 96 well plates and inoculated with compounds with each inhibitor, along with DMSO or RIF controls. The luciferase assay was carried out as previously described and plates were read on a PerkinElmer Enspire plate reader.

For studies in macrophages, primary bone marrow derived macrophages were harvested and infected as previously described ^48^. Briefly, BMMΦ from C57Bl/6 mice were distributed into 96 well white plates and infected for 1 hour with a CDC1551 luciferase reporter strain^47^. Following 1 hour of infection, cells were treated with inhibitors ranging from 200 – 0.2 μM of MmpL3 inhibitors. 20 μM PAS, 3 μM RIF, and DMSO were used as controls. Samples were incubated in the 96 wells plates at 37°C + 5% CO_2_ for 6 days before carrying monitoring bacterial survival by measuring luciferase activity. Experiments were conducted in biological triplicate and repeated at least once with similar results.

### Protein Modeling

The 3D structure for MmpL3 was generated using the I-TASSER server^49^. The MmpL3 protein sequence of H37Rv from Mycobrowser (Rv0206c) ^50^ was aligned to the MmpL3 crystal structure of Msm (PDB: 6AJF) with a resulting C-score of 0.17 (TM-score 0.74 ± 0.11, RMSD 8.4 ± 4.5Å). The resulting structure was modified to remove the C-terminal tail (732/944AA) in PyMol 2.2.3 ^51^.

### DiaMOND

DiaMOND analysis was carried as described by Cokol *et al.*, with modifications as described (12). Briefly concentration ranges were linearized using the equation

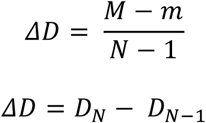

Where *ΔD* is the difference between concentrations of each dose, *M* is the lowest concentration to inhibit Mtb growth 100%, and *m* is the highest concentration estimated to confer 0% growth inhibition based on the EC_50_ dose response curves. *N* is the number of doses to be used in DiaMOND. Mtb was then treated with each concentration range for each compound by itself (Null treatment) at a [X_N_] or in combination with another inhibitor at a [½X_N_]. Dose responses were used to generate a dose response curve for each treatment which was used to interpolate the IC_50_ which was set for the observed, “*o*” to calculate the 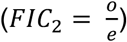 as previously described^37^. Dose responses were conducted in biological duplicate and reported FIC_2_ values are representative of the geometric mean of two reps.

### Eukaryotic Cytotoxicity

Primary BMMΦ were isolated and distributed into white wall 96 well plates as described above. Cells were treated with inhibitors ranging in concentration from 200 – 0.26 μM. Cells were incubated at 37C for 3 or 6 days with 5% CO_2_. Cytotoxicity was tested using the Cell Titer Glow assay kit using the methods from the provider. For a negative control cells were treated with 4% of TritonX-100 and DMSO as a positive control ^28^.

### Kinetic solubility and microsomal stability assay

The kinetic solubility assay was conducted as described by Bevan *et al.*^*52*^ Briefly, the assay was performed with 7-point (2-fold) dilutions from 200 μM - 3.125 μM for the compounds. Mebendazole, benxarotene and aspirin were also included as controls. The drug dilutions were added to PBS, pH 7.4, with the final DMSO concentration of 1%, and incubated at 37 °C for 2 h. The absorbance at 620 nm was measured for each drug dilution to estimate of the compound solubility. Three replicates were examined for each dilution. Mouse microsomal stability was conducted as described by Obach ^53^ and presented as % remaining following 30 minutes. Values greater than 100% are likely due to changes in the solubility of the compounds over the course of the assay and represent high stability in microsomes.

## Acknowledgements

We thank Bree Aldridge for assistance in designing the DiaMOND studies. Christopher Colvin and Benjamin Johnson provided technical assistance in the initial prioritization of compounds. Additional technical assistance was provided by Bilal Alewei and Tom Dexheimer, in the MSU medicinal chemistry and screening cores to conduct the solubility and microsomal stability studies. Research reported in this study was supported by start-up funding from Michigan State University and AgBioResearch, and grants from the NIH-NIAID (U54AI057153, R21 AI1170181 and R01AI116605), the Bill and Melinda Gates Foundation (OPP1059227), a strategic partnership grant from the MSU Foundation (14-SPG-Full-2966), and the Jean P. Schultz Endowed Biomedical Research Fund at the MSU College of Human Medicine.

## Author Contributions

JW and RBA designed the experiments and wrote the manuscript; GC conducted prioritization assays; JW, EH, KC conducted the Mtb and spectrum of activity experiments; CC and TD designed and conducted the *M. abscessus* experiments; and, EE directed the solubility and microsomal stability studies. All authors reviewed the manuscript.

## Supplemental Methods for *M. abscessus* spectrum of activity assays

### Bacterial strains and culture media

For screens and hit confirmation, *Mycobacterium abscessus* Bamboo was used. *M. abscessus* Bamboo was isolated from the sputum of a patient with amyotrophic lateral sclerosis and bronchiectasis and was provided by Wei Chang Huang, Taichung Veterans General Hospital, Taichung, Taiwan. *M. abscessus* Bamboo whole genome sequencing showed that the strains belongs to *M. abscessus* subsp. *abscessus* and harbors an inactive clarithromycin-sensitive *erm* C28 sequevar (GenBank accession no. <u>MVDX00000000</u>). *M. abscessus* Bamboo cultures were grown in standard mycobacterium medium, Middlebrook 7H9 broth (BD Difco) supplemented with 0.5% albumin, 0.2% glucose, 0.085% sodium chloride, 0.0003% catalase, 0.2% glycerol, and 0.05% Tween 80. Solid cultures were grown on Middlebrook 7H10 agar (BD Difco) supplemented with 0.5% albumin, 0.2% glucose, 0.085% sodium chloride, 0.5% glycerol, 0.0003% catalase, and 0.006% oleic acid.

### Single-point growth inhibition screening assay

The compound library was screened in microtiter plates as previously described with minor modifications. Briefly, the screen was carried out in 96-well flat-bottom Corning Costar plates at a single-point concentration of 20 μM with a starting inoculum of an optical density at 600 nm (OD_600_) of 0.05 (10^7^ CFU/ml) in a final volume of 200 μl. The culture for the starting inoculum was diluted from a preculture at mid-log phase (OD_600_, 0.4 to 0.6). The plates were sealed using a Breathe-Easy sealing membrane (Sigma-Aldrich), put in an airtight container with moist tissue, and incubated for 3 days at 37°C on an orbital shaker at 110 rpm. Each plate had a medium-only control and a drug-free control, as well as positive control, clarithromycin at 20 μM. After 3 days of incubation, the cultures in the wells were manually resuspended before the OD_600_ was read in a TECAN Infinite Pro 200 plate reader. Compounds were scored according to their growth inhibition of the treated culture compared to the untreated culture (DMSO-treated). The experiment was conducted in duplicate, and the results are shown as a scatter plot, with each data point representing the mean of data from the two replicates for each compound (Fig. 1).

### Growth inhibition dose-response assay

MICs in dose-response assays were determined by the broth microdilution method as described previously (63), with some modifications. Briefly, 96-well plates were filled with 100 μl of 7H9 medium in each well. Two times the desired two-fold (10 points) serial dilutions of compounds were prepared with TECAN D300e Digital Dispenser. An appropriate dilution of a mid-log-phase culture to an OD_600_ of 0.1 (final OD_600_ in all wells was 0.05) was carried out, and 100 μl of the bacterial culture was added to the wells. The plates were incubated at 37°C and 110 rpm on an orbital shaker for 3 days and then manually resuspended, and the OD_600_ was measured using the plate reader. We report MIC_50_s and MIC_90_s which are the concentrations that inhibit 50% and 90% of growth respectively compared to the untreated control. The MIC_90_s correspond to the standard “no visible growth” MICs. All experiments were carried with biological replicates as well as technical replicates.

## Supplemental Figure Legends

**Figure S1:**
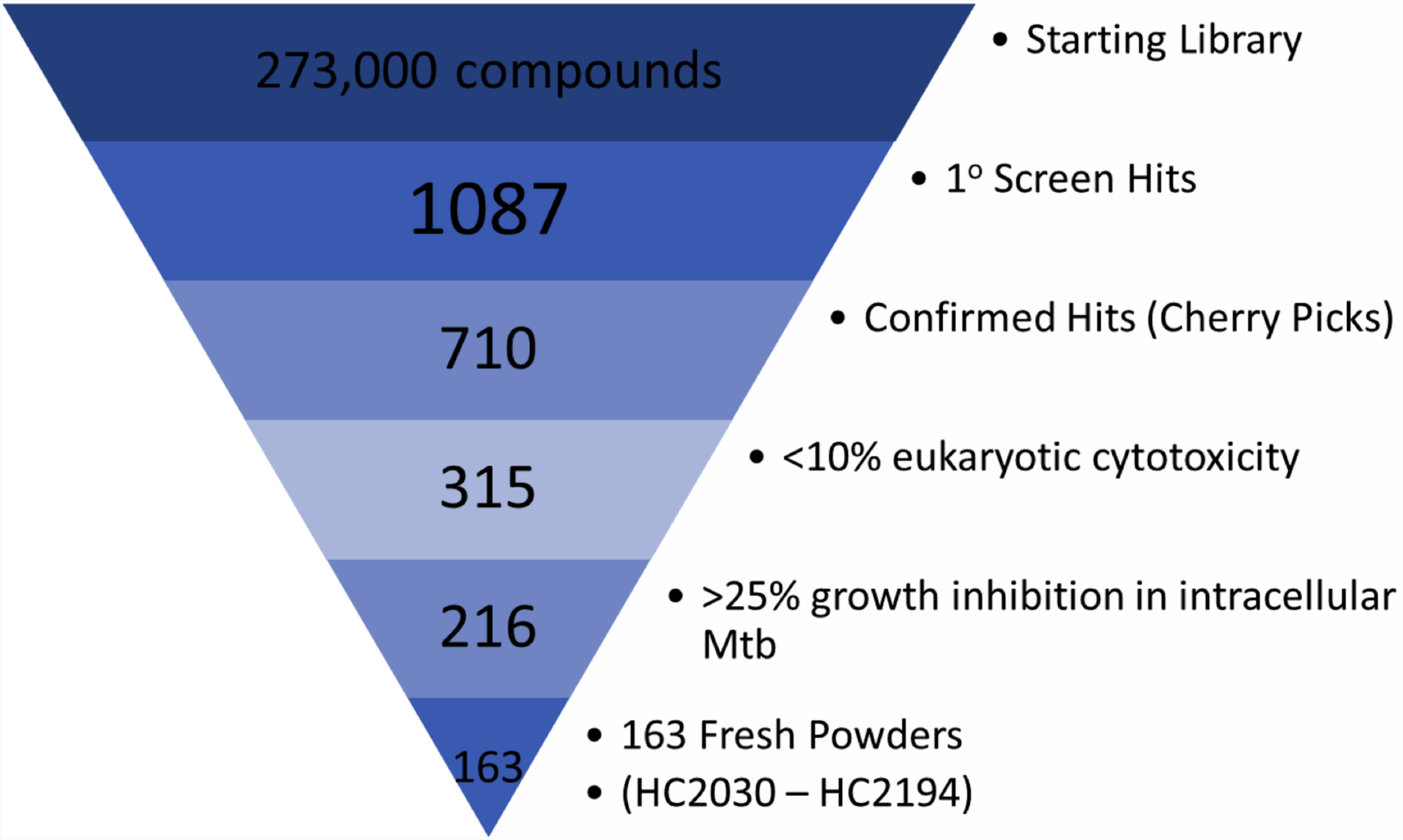
Prioritization funnel of growth inhibitors from a high throughput screen. A high throughput screen of 273,000 compounds identified 1087 compounds that inhibit Mtb growth independent of the targeted two component regulators at 10 μM. These compounds were further tested as being able to inhibit Mtb growth (confirmed hits), have low eukaryotic cytotoxicity (<10%), able to inhibit intracellular Mtb growth (>25%) resulting in 216 compounds that meet the minimum requirements. Of the 216 compounds 163 commercially available compounds were purchased as fresh powders.

**Figure S2:**
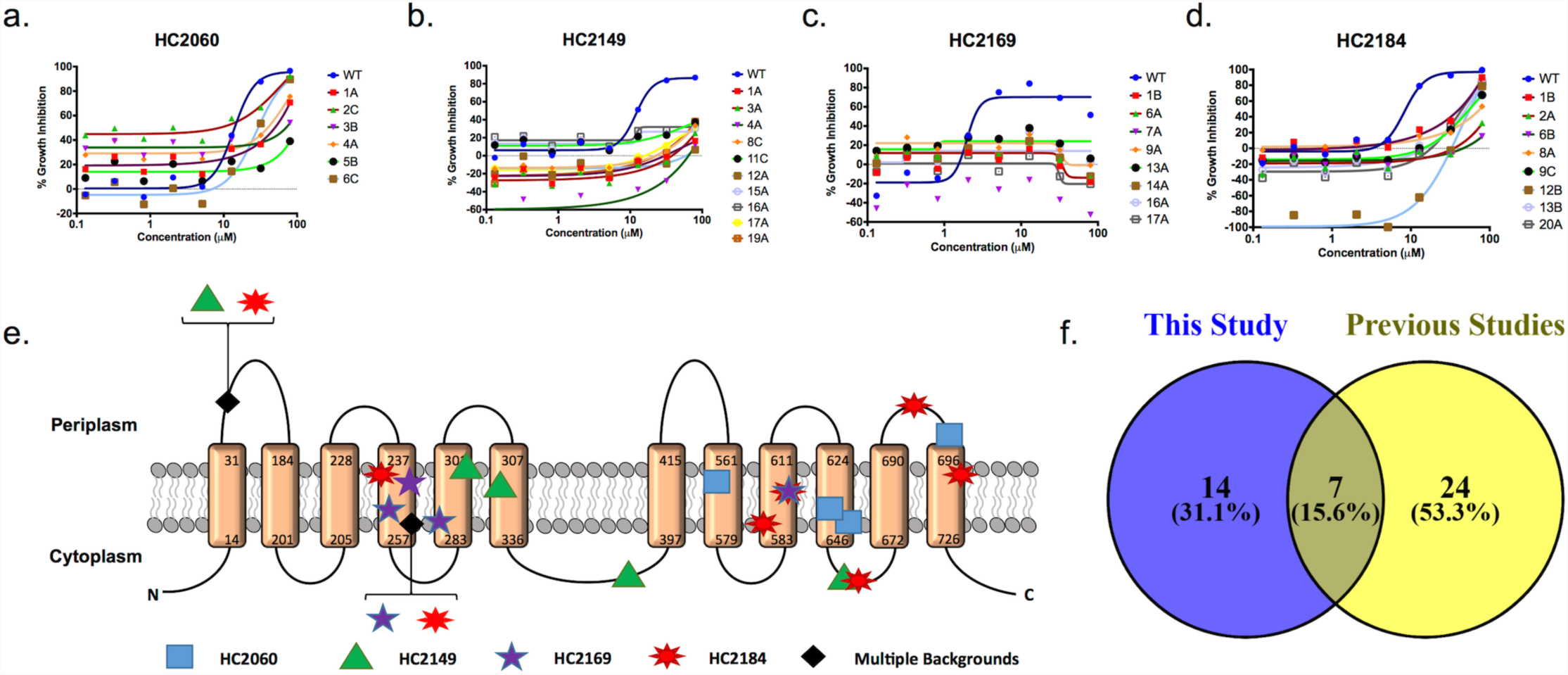
Resistant Mutants to Four Novel Inhibitors Map to mmpL3. a-d) Dose response curves of resistance mutant to four novel Mtb growth inhibitors. Curves are based on 2.5 dilutions of inhibitors ranging from 80 to 0.13μM. Experiments were conducted in triplicate and the error bars indicated the standard deviation of the mean. e) Transmembrane domain map shows diversity of substitutions conferred by mutations in mmpL3. Transmembrane domain map is based on Phyre2 analysis of H37Rv MmpL3 protein sequence. f) Venn Diagram identifies novel MmpL3 substitutions identified in this study. A total of 21 MmpL3 amino acid substitutions were identified in this study, including 14 novel substitutions and 7 previously identified substitutions (see TABLE S1 for list of substations).

**Figure S3.**
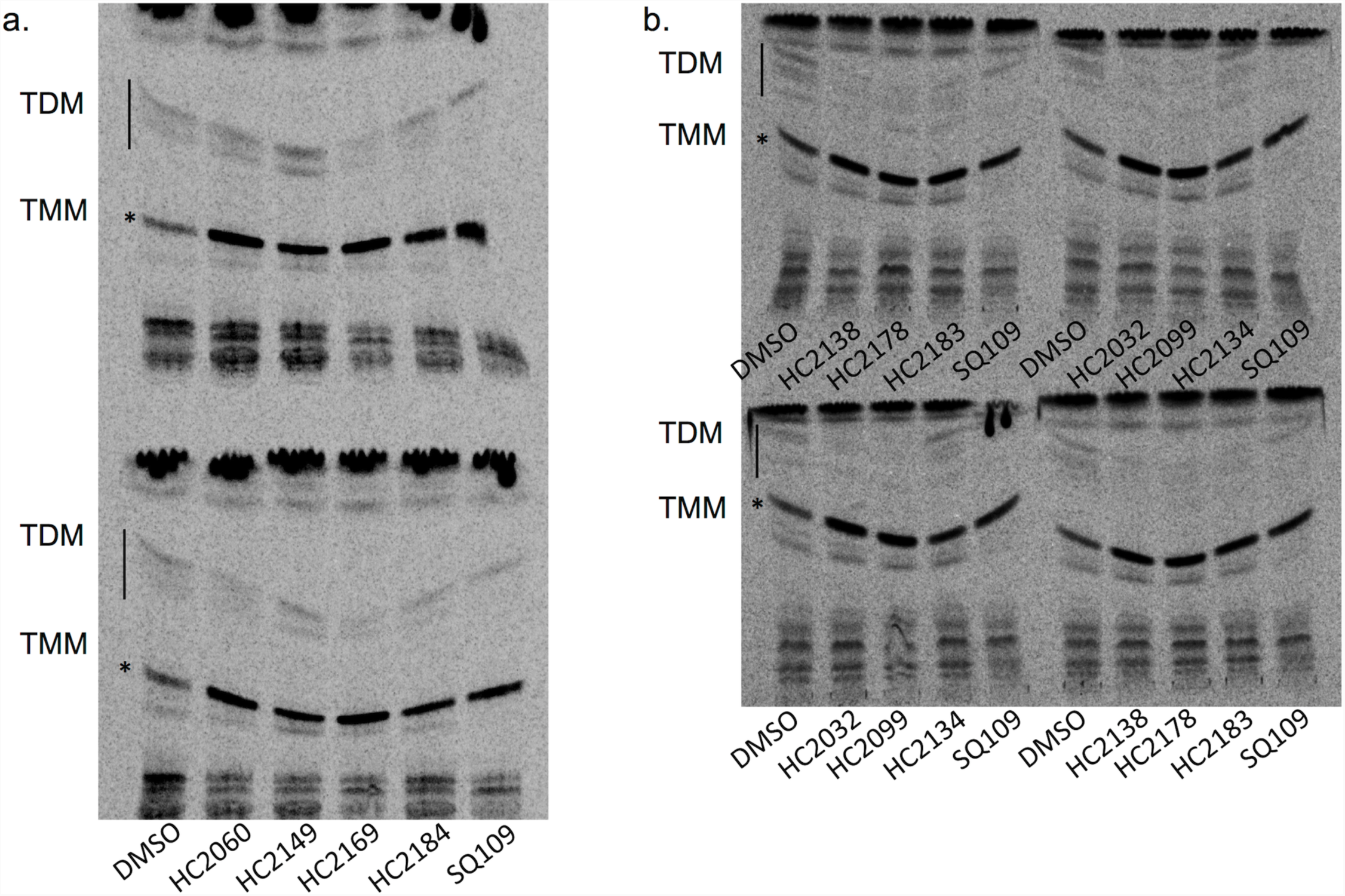
TLCs show TMM/TDM Modulation. **a-c)** Mtb cells were grown in the presence of 8μCi of 14C-acetate for twenty four hours and treated with 20μM of **a)** the four prioritized inhibitors (HC2060, HC2149, HC2169 or HC2184) or **b and c)** the six inhibitors identified from the targeted mutant phenotypic screen (HC2032, HC2099, HC2134, HC2138, HC2178, HC2183). Lipids were isolated from whole cell extracts and analyzed by TLC. In each experiment samples of cells were also treated with either 20μM of SQ109 or DMSO.

**Figure S4.**
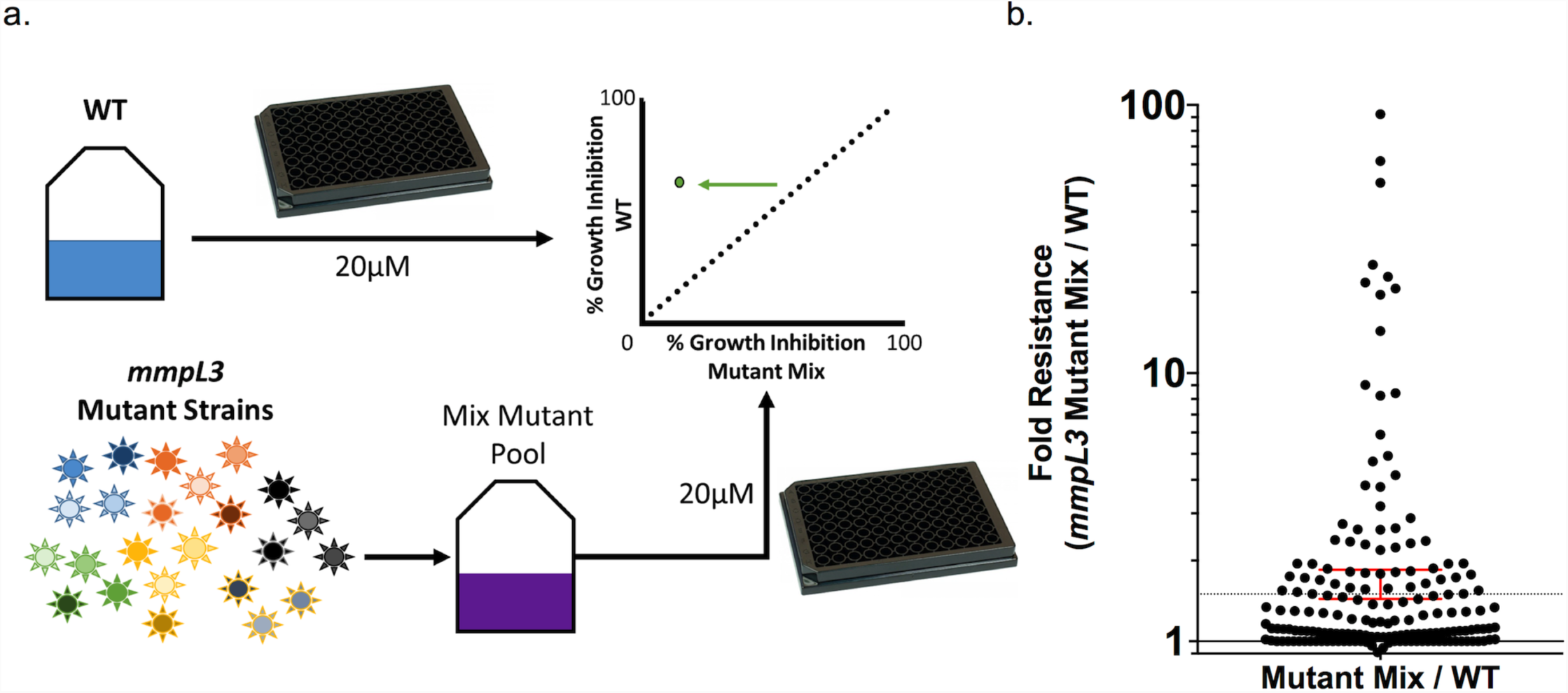
Illustrated Outline of Targeted Mutant Phenotypic Screen. **a)** Growth inhibition of a pooled culture of twenty four unique *mmpL3* mutant strains of Mtb (multicolored suns) is directly compared with WT Mtb strains. Samples of either pooled *mmpL3* mutant strains or WT Mtb are aliquoted into separate 96 well plates and treated with 163 prioritized Mtb growth inhibitors, as well as BDQ, CFZ, INH, PAS, SQ109 and H_2_O_2_. Growth inhibition (%) is calculated as the growth inhibition relative to the DMSO and RIF controls. **b)** Beehive plot of relative fold decrease in activity of compounds in the mixed *mmpL3* mutant background compared to WT treated cells. Dotted line indicates a 1.5 fold resistance in the *mmpL3* mixed mutant background relative to the WT. Error bars (red) indicate the 95% confidence interval of the geometric mean.

**Figure S5.**
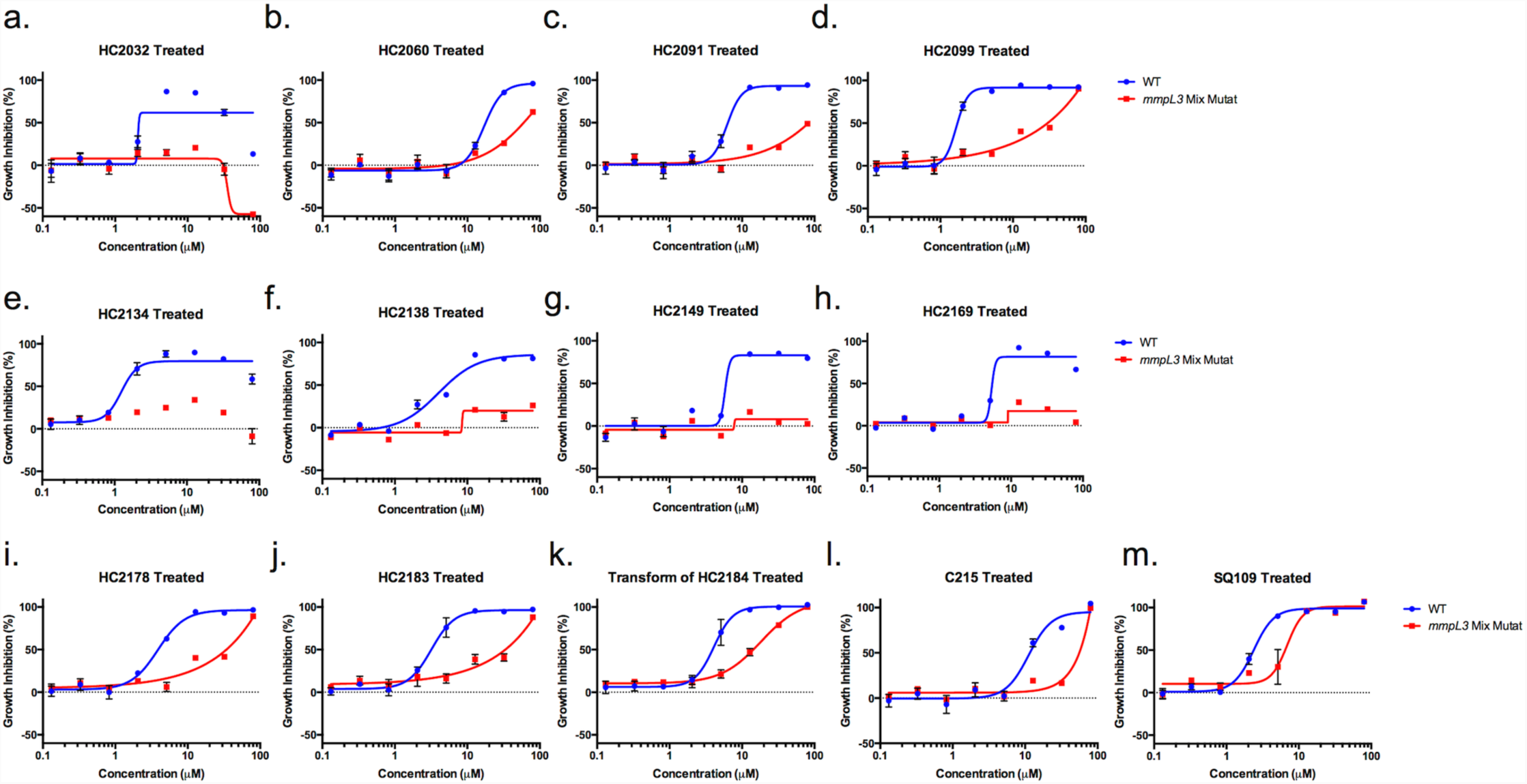
13 Dose response curves of thirteen proposed MmpL3 inhibitors on pooled mmpL3 mutant strains. **a-m)** Dose response curves of thirteen Mtb growth inhibitors confirmed to have reduced activity in the pooled mmpL3 mutant background (red) compared to WT Mtb (blue). Samples were treated with a series of (2.5 fold dilutions) of each inhibitor ranging form 80μM to 0.13μM. Growth inhibition (%) is calculated as the growth inhibition relative to the DMSO and RIF controls. Experiments were conducted in triplicate and error bars indicate the standard deviation from the mean.

**Figure S6.**
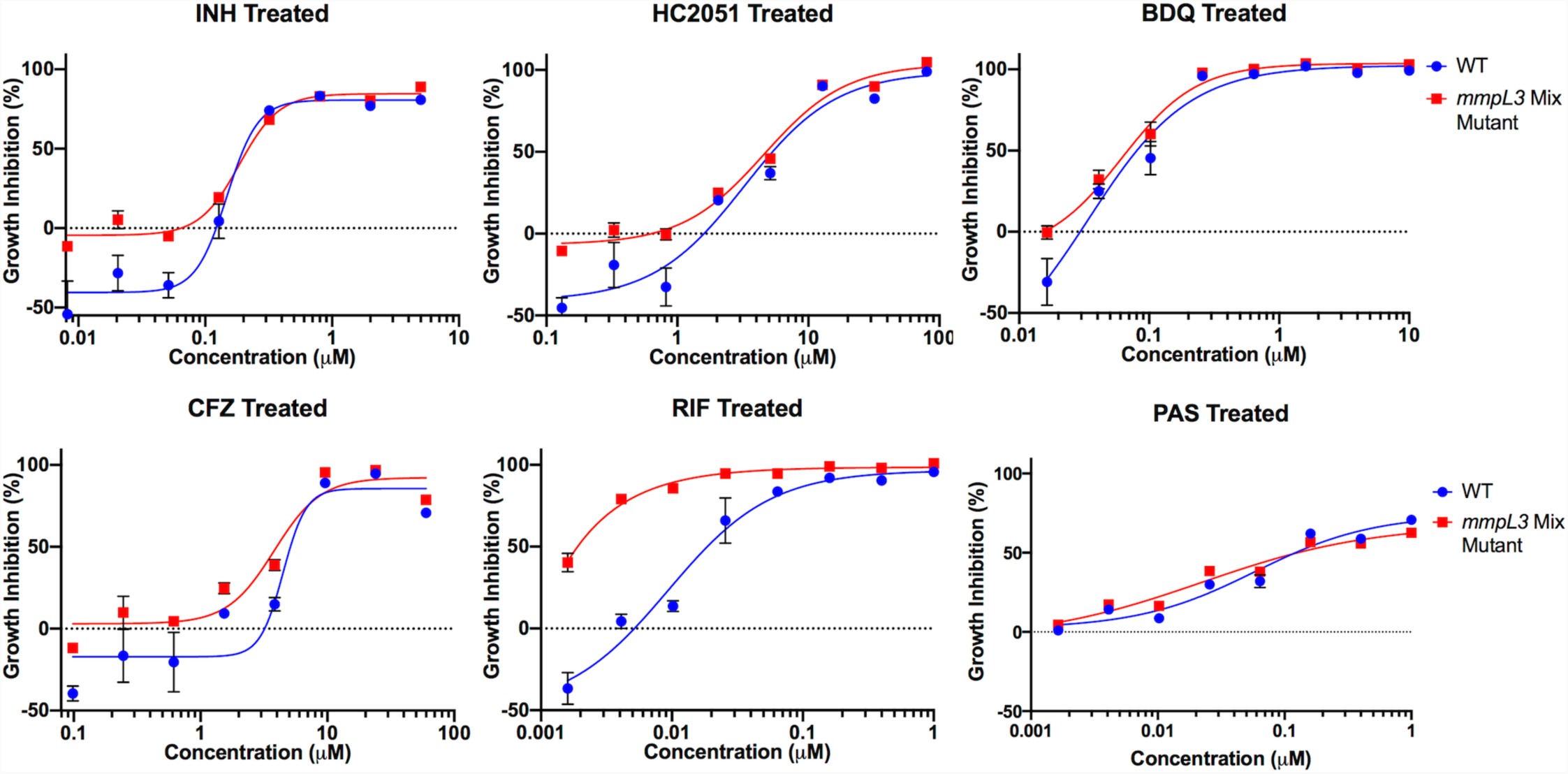
Impact of Non-MmpL3 inhibitors on pooled mmpL3 mutant strains. Dose response curves of thirteen Mtb growth inhibitors confirmed to have reduced activity in the pooled mmpL3 mutant background (red) compared to WT Mtb (blue). Samples were treated with a series of (2.5 fold dilutions) of each inhibitor ranging form 80μM to 0.13μM. Growth inhibition (%) is calculated as the growth inhibition relative to the DMSO and RIF controls. Samples were run in triplicate and error bars indicate the standard deviation from the mean.

**Figure S7.**
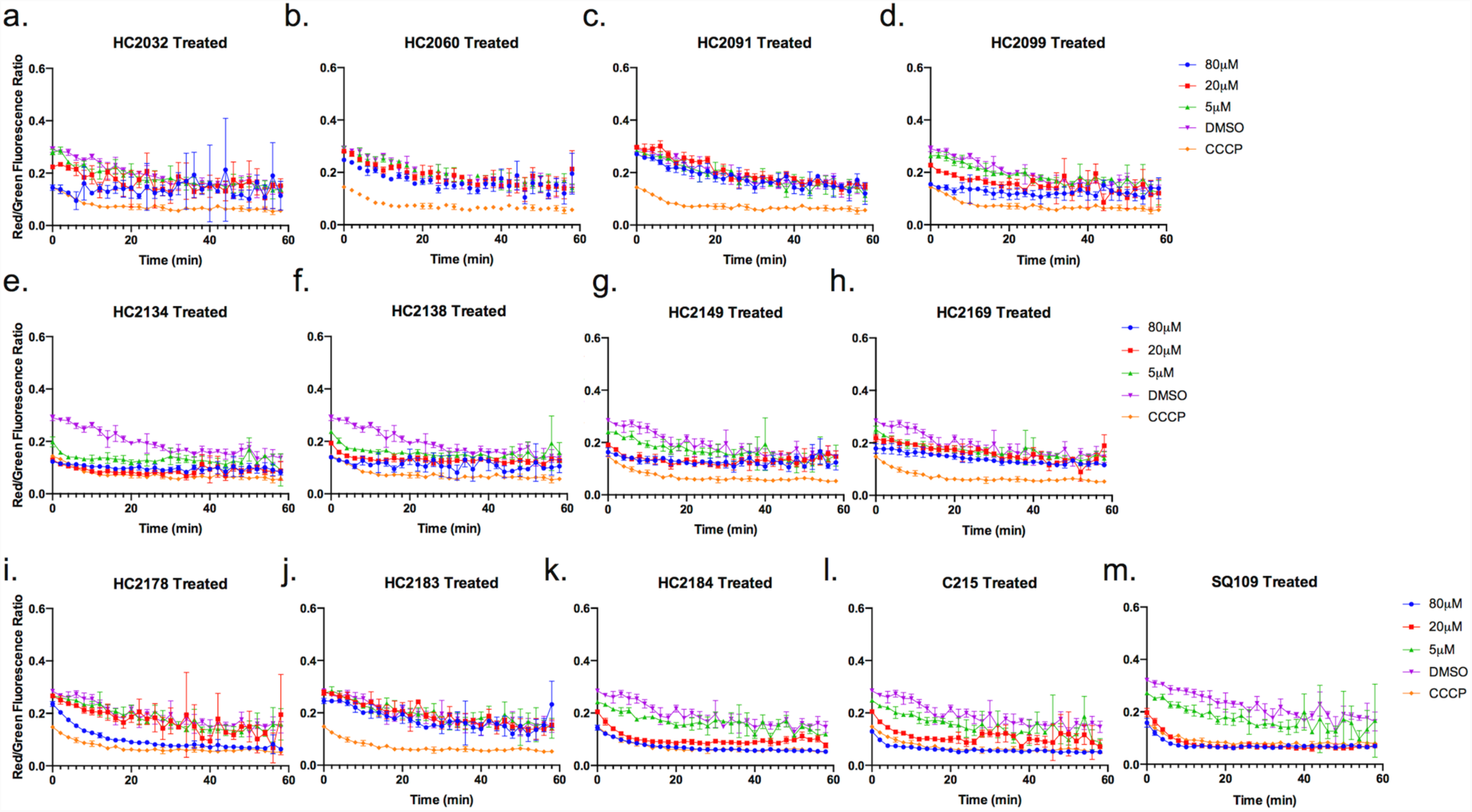
Capacity of Inhibitors to Disrupt Membrane Potential. **a-m)** Mtb cells labeled with DiOC2 and treated with 80μM (blue circle), 20μM (red square) or 5μM (green triangle) of each of the thirteen MmpL3 inhibitors for one hour. As controls DMSO (negative, purple inverted triangles) and CCCP (positive, orange diamonds) treatments were also included. Experiments were carried out using the DiOC_2_ membrane potential assay kit. The experiment was repeated twice with similar results. Data points are the geometric mean of three technical repeats. Error bars indicate the geometric standard deviation of three technical replicates. The experiment was repeated with similar results.

**Figure S8.**
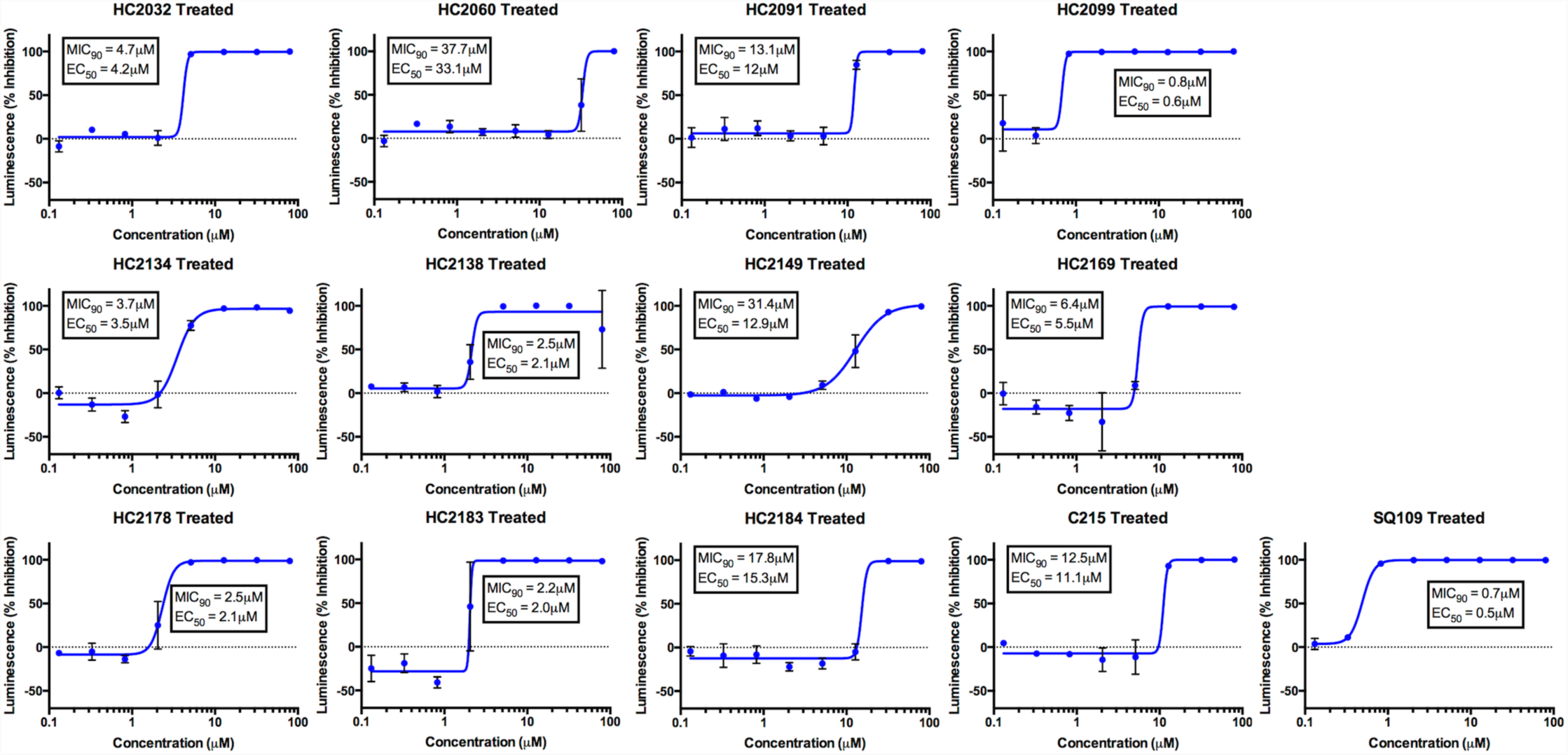
Bactericidal activity of MmpL3 Inhibitors. Mtb CDC1551 luc reporter strains were treated with a series of dilutions (2.5 fold) from 80 to 0.13μM of each of the thirteen MmpL3 inhibitors for six days in vitro. Cells were then tested for luciferase expression using the Bright-Glo Luciferase assay kit. Growth inhibition (%) is the normalized luciferase activity relative to the DMSO – positive and RIF – negative controls. Experiments were conducted in triplicate and the error bars indicated the standard deviation of the mean.

**Figure S9.**
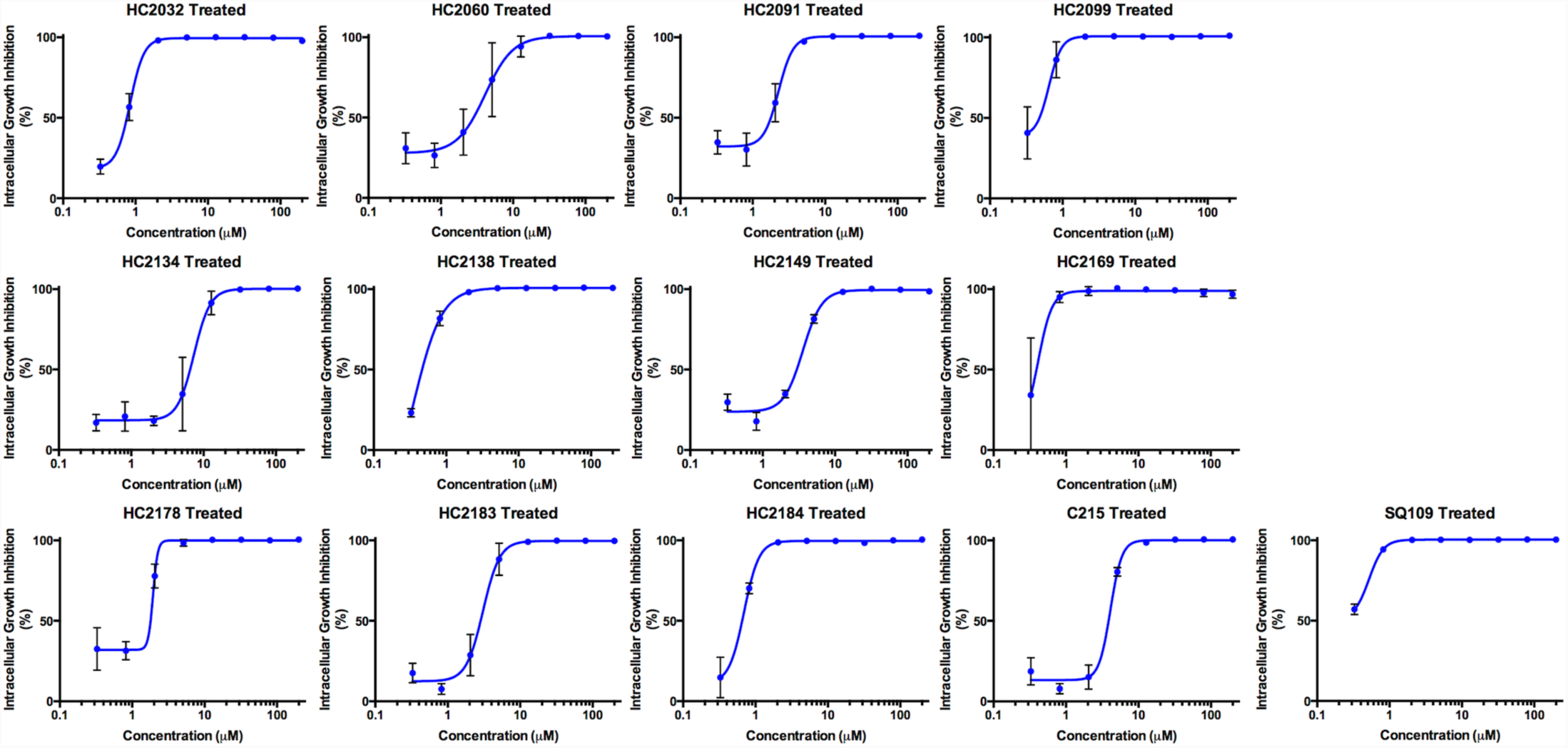
Impact of MmpL3 Inhibitors on Intracellular Growth. Primary bone marrow macrophages were infected with Mtb CDC1551 luc reporter strains. Infected macrophages were treated with a series of dilutions (2.5 fold) from 200 to 0.3μM of each of the thirteen MmpL3 inhibitors for six days in vitro. Cells were then tested for luciferase expression using the Bright-Glo Luciferase assay kit. Growth inhibition (%) is the normalized luciferase activity relative to the DMSO – positive and RIF – negative controls. Experiments were conducted in triplicate and the error bars indicated the standard deviation of the mean.

**Figure S10.**
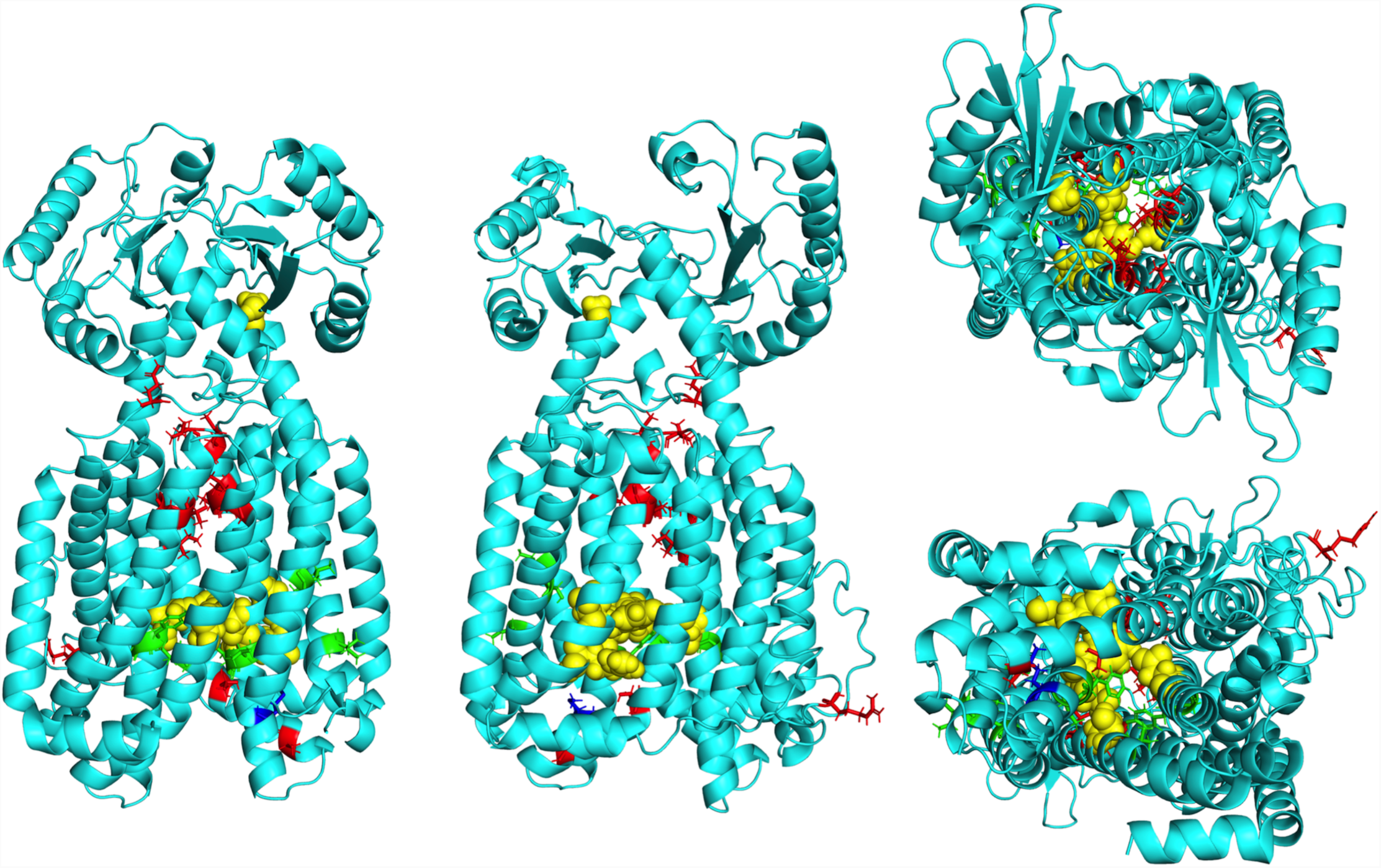
Clade substitutions Differ in Proximity to Essential Residues. **a-d)** Front, back, top, and bottom (respectively) views of an I-TASSER predicted structure of Mtb MmpL3 based on MmpL3 structure of Msm (PDB: 6AJH). Substitutions conferred by mutations in *mmpL3*. Substitutions are colored based on clade from cross resistance profiling, Clade I substitutions (green), Clade II substitutions (red), or M649 (blue) which fell into both clades depending on substitution. Yellow spheres indicate the seven essential residues (D251, S288, G543, D640, Y641, D710, and R715) for MmpL3 activity identified by Bellardinelli and colleagues^17^. The model shows a truncated version (732/944aa) of the MmpL3 protein lacking the C-terminal tail.

**TABLE S1.** Sequencing Results of Resistant Mutants

| Compound | Strain | SNP Location (nt) | Quality Score | Gene | Nucleotide Change | AA Change |
| --- | --- | --- | --- | --- | --- | --- |
| **HC2060** | 1A / 2C | 245506 | 5308 / 4597 | *mmpL3* | GTG --> **ATG** | V643M |
|  | 3B | 245501 | 4710 | *mmpL3* | TTC --> TT**G** | F644L |
|  | 4A | 245733 | 3840 | *mmpL3* | CTG --> C**CG** | L567P |
|  | 5B | 245501 | 4807 | *mmpL3* | TTC --> TT**A** | F644N* |
|  | 6C | 245349 | 4795 | *mmpL3* | ATG --> A**CG** | M695T* |
| **HC2149** | 1A | 245487 | 5798 | *mmpL3* | ATG --> A**CG** | M649T* |
|  | 3A / 4A / 8C / 17A / 19A | 247313 | 5895 / 6057 / 5228 / 4998 / 5725 | *mmpL3* | CAG --> CA**T** | Q40H* |
|  | 11C / 12A | 246316 | 5127 / 6751 | *mmpL3* | CGG --> T**GG** | R373W* |
|  | 15A | 246501 | 3836 | *mmpL3* | ACC --> A**TC** | T311I* |
|  | 16A | 246537 | 3854 | *mmpL3* | CTG --> C**AG** | L299Q* |
| **HC2169** | 1B / 6A / 17A | 245662 | 7164 / 6152 / 6805 | *mmpL3* | TCG --> **ACG** | S591T* |
|  | 14A | 246579 | 4739 | *mmpL3* | GTG --> G**GG** | V285G* |
|  | 13A | 246675 | 4585 | *mmpL3* | GGG --> G**AG** | G253E |
|  | 7A / 9A | 246678 | 6076 / 4312 | *mmpL3* | TAC --> T**GC** | Y252C* |
|  | 16A | 246702 | 5280 | *mmpL3* | ATC --> A**CC** | I244T* |
| **HC2184** | 1B | 245355 | 5122 | *mmpL3* | GAC --> G**GC** | L693P* |
|  | 2A | 246675 | 6093 | *mmpL3* | CCC --> C**TC** | G253E |
|  | 6B | 245661 | 5719 | *mmpL3* | TCG --> T**AG** | S591I |
|  | 8A | 246678 | 5654 | *mmpL3* | ATG --> A**CG** | I585S* |
|  | 9C | 247313 | 5824 | *mmpL3* | GTC --> GT**G** | Q40H* |
|  | 12B | 245338 | 5281 | *mmpL3* | GAC --> T**AC** | L699M |
|  | 13B | 245448 | 5121 | *mmpL3* | CGC --> C**TC** | A662E* |
|  | 20A | 246714 | 3800 | *mmpL3* | CAC--> C**GC** | V240A* |

\* - Novel substitutions not previously identified in Mtb

**TABLE S2.**
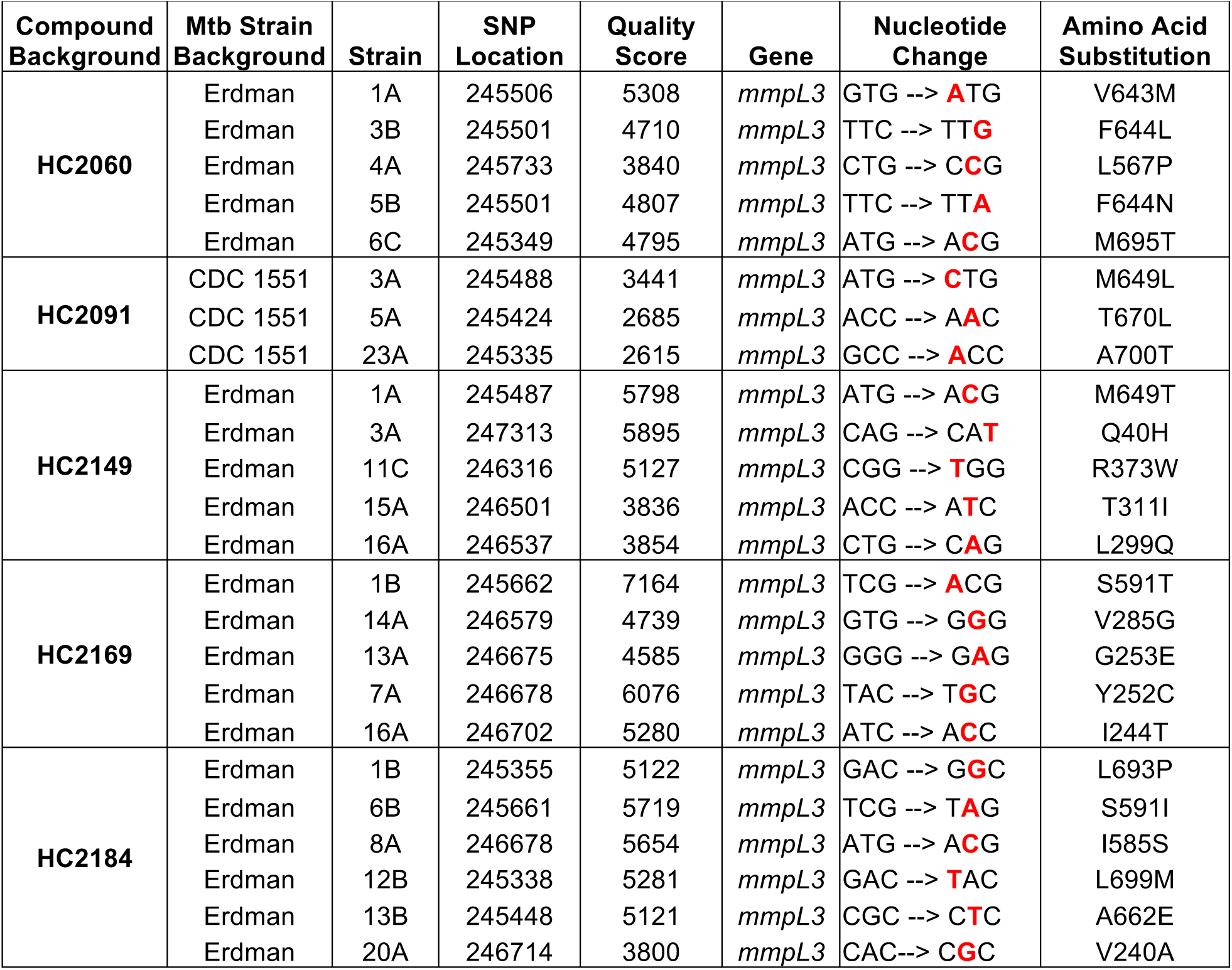
Genetic background of Mtb strains used in Screen

**TABLE S3.** EC_50_ Values of Control Compounds

| Treatment | WT EC<sub>50</sub> | Mix Mutant EC<sub>50</sub> | 95% Confidence Interval |
| --- | --- | --- | --- |
| **INH** | 0.15 | 0.18 | n.s. |
| **HC2051** | 3.2 | 4.5 | n.s. |
| **CFZ** | 4.4 | 3.8 | n.s. |
| **BDQ** | 0.03 | 0.06 | n.s. |
| **RIF** | 0.009 | <0.009 | N.D. |
| **PAS** | 0.05 | 0.02 | n.s. |

n.s. – Not significant
N.D. – Not determined

**Table S4.** AUC Values from Cross Resistance Profiling

|  | HC2032 | HC2060 | HC2091 | HC2099 | HC2134 | HC2138 | HC2149 | HC2169 | HC2178 | HC2183 | HC2184 | C215 | SQ109 |
| --- | --- | --- | --- | --- | --- | --- | --- | --- | --- | --- | --- | --- | --- |
| Q40H<sup>E</sup> | 42.98 | 68.03 | 84.3 | 101.8 | 63.23 | 37.49 | 44.07 | 42.17 | 92.35 | 96.53 | 63.86 | 131.7 | 143 |
| V240A<sup>E</sup> | 102.2 | 82.81 | 73.08 | 89.43 | 68.44 | 15.86 | 77.88 | 94.26 | 108.7 | 107.7 | 68.1 | 26.77 | 143.3 |
| I244T<sup>E</sup> | 110.9 | 69.62 | 60.8 | 67.29 | 86.09 | 65.59 | 58.91 | 52.76 | 97.73 | 96.78 | 73.2 | 33.41 | 132.8 |
| T252C<sup>E</sup> | 149.4 | 48.76 | 27.29 | 55.58 | 35.23 | 14.64 | 23.38 | 35.96 | 87.77 | 60.9 | 58.37 | 18.72 | 181.4 |
| G253E<sup>E</sup> | 18.23 | 60.32 | 18.2 | 44.63 | 11.15 | 7.195 | 10.61 | 29.13 | 54.42 | 57.9 | 32.29 | 15.03 | 146 |
| V285G<sup>E</sup> | 31.87 | 40.62 | 39.38 | 48.43 | 29.35 | 34.77 | 34.81 | 41.29 | 78.09 | 57.41 | 59.12 | 33.96 | 117.1 |
| L299Q<sup>E</sup> | 95.93 | 85.39 | 72.67 | 90.14 | 59.81 | 37.88 | 21.73 | 55.48 | 91.53 | 93.22 | 73.78 | 40.02 | 157.2 |
| T311I<sup>E</sup> | 94.83 | 84.38 | 101.8 | 115.7 | 106.2 | 68.61 | 23.31 | 54.41 | 114.1 | 123.6 | 96.01 | 127.8 | 167.8 |
| R373W<sup>E</sup> | 62.68 | 78.36 | 70.65 | 79.96 | 53.12 | 85.28 | 37.85 | 35.57 | 105.8 | 91.78 | 100.1 | 62.15 | 157.2 |
| L567P<sup>E</sup> | 43.81 | 40.82 | 40.23 | 52.93 | 35.96 | 69.17 | 120.2 | 157.9 | 80.5 | 61.52 | 69.83 | 137.5 | 157.3 |
| I585S<sup>E</sup> | 165.1 | 47.59 | 26.81 | 40.55 | 31.87 | 53.06 | 23.38 | 19.76 | 90.54 | 34.68 | 35.16 | 31.11 | 201.3 |
| S591I<sup>E</sup> | 28.78 | 21.79 | 76.97 | 38.94 | 18.47 | 18.89 | 12.86 | 22.95 | 35.47 | 55.81 | 28.15 | 11.45 | 110.7 |
| S591T<sup>E</sup> | 31.85 | 39.27 | 66.47 | 65.59 | 68.16 | 43.98 | 24.29 | 28.08 | 80.44 | 90.53 | 50.04 | 32.04 | 144.8 |
| V643M<sup>E</sup> | 33.28 | 40.88 | 28.28 | 41.06 | 36.34 | 48.63 | 38.99 | 98.7 | 56.54 | 46.87 | 54.33 | 88.44 | 117.9 |
| F644L<sup>E</sup> | 34.12 | 21.31 | 25.91 | 48.32 | 130.9 | 123.7 | 93.34 | 58.46 | 51.75 | 38.31 | 65.11 | 48.58 | 132.2 |
| F644N<sup>E</sup> | 25.76 | 27.24 | 38.88 | 53.13 | 123.5 | 101.5 | 95.12 | 54.82 | 54.16 | 54.71 | 82.96 | 14.76 | 117.6 |
| M649L<sup>C</sup> | 22.92 | 82.01 | 38.01 | 68.54 | 73.58 | 72.83 | 47.07 | 28.83 | 96.72 | 41.98 | 62.24 | 55.8 | 175.5 |
| M649T<sup>E</sup> | 43.61 | 72.15 | 47.2 | 66.99 | 84 | 82.72 | 17.65 | 23.53 | 105.5 | 97.45 | 67.32 | 50.67 | 171.8 |
| A662E<sup>E</sup> | 100.1 | 101.7 | 51.92 | 83.82 | 132.1 | 129.6 | 69.05 | 44.16 | 134.3 | 110.9 | 70.83 | 37.18 | 183.2 |
| T670L<sup>C</sup> | 75.25 | 75.14 | 21.15 | 52.09 | 109.1 | 118.3 | 56.69 | 71.4 | 151.6 | 64.69 | 59.74 | 88.44 | 174.2 |
| L693P<sup>E</sup> | 88.03 | 73.82 | 86.89 | 88.84 | 96.19 | 107.3 | 83.23 | 39.29 | 93.76 | 111.5 | 63.13 | 71.82 | 169.4 |
| M695T<sup>E</sup> | 79.44 | 55.82 | 89.55 | 103.1 | 92.69 | 45.96 | 61.88 | 79.04 | 60.96 | 103.9 | 68.95 | 92.96 | 139.8 |
| L699M<sup>E</sup> | 55.85 | 69.13 | 57.75 | 89.07 | 90.94 | 42.28 | 60.88 | 69.08 | 90.47 | 90.9 | 66 | 51.01 | 134.2 |
| A700T<sup>C</sup> | 69.86 | 79.3 | 43.85 | 78.75 | 90.92 | 132.3 | 59.37 | 45.64 | 106.9 | 99.82 | 83.75 | 61.3 | 151.5 |
| WT-CDC1551 | 136.2 | 93.86 | 114 | 129.8 | 159 | 150.7 | 96.38 | 151.5 | 148.2 | 154.7 | 123 | 97.66 | 193.8 |
| Average<br>WT -<br>Erdman | 115.96 | 74.87 | 101.18 | 115.1 | 142.74 | 140.3 | 89.33 | 141.14 | 130.32 | 138.96 | 94.88 | 69.12 | 171.88 |

E or C indicates the WT background that the AUC was compared to. E – Erdman and C – CDC1551

